# The Genetic Landscape of Structural Variants in *Aspergillus flavus* using ONT and PacBio Sequencing

**DOI:** 10.64898/2026.10.05.756768

**Authors:** Alexie Woods, Jessica M. Lohmar, Amna Malik, Arthur J. Lustig

## Abstract

Genetic rearrangement in the absence of selection is poorly understood. Long-read, high-depth sequencing has enabled analysis of the genetic landscape of spontaneous structural variants. We have applied PacBio Revio sequencing to identify the major classes of genetic alterations during long-term vegetative growth in *Aspergillus flavus*. These classes include a) precise deletions frequently associated with the flanking region of homology, b) clusters of short deletions that are heavily enriched within centromeric and subtelomeric regions that are typically 94% A+T, and c) deletions and expansions within a subset of homopolymer tracts. We find 70 homopolymer tracts longer than 44 bp, dispersed throughout the genome in noncoding regions and unassociated with transposable elements. A broader analysis of third-generation sequenced datasets from fungi found within the Phylum Ascomycota showed a high but variable prevalence of homopolymeric tracts across species. The adventitious identification of a putative *flbA* mutation, responsible for an altered conidial phenotype, by the Deep Variant AI system also demonstrates the power of this approach. These studies show that high-depth, long-read sequencing can probe SNPs, indels, homopolymeric tracts, and large-scale rearrangements in an unbiased fashion, and demonstrate that short deletions and homopolymer tract expansions arise spontaneously in *A. flavus* and other filamentous fungi.

## INTRODUCTION

Genetic instability that occurs in the absence of selection underpins our fundamental understanding of genomic rearrangements and disease states (Brown and Freudenreich 2021). Spontaneous recombination has long been studied as a means of understanding specific classes or sites of recombination; however, an unbiased assessment of such recombination events has been challenging (Jinks-Robertson and Petes 2021). These events are difficult to study using standard genetic techniques, owing to their low frequency and the difficulty in identifying complex forms (Reams and Roth 2015).

Several model systems and technological advances have helped address this limitation. In the yeasts *Saccharomyces cerevisiae* and *Schizosaccharomyces pombe,* the most common genetic changes are single nucleotide polymorphisms (SNPs), regions with loss of heterozygosity (LOH), and small insertions and deletions (indels) (Yang Sui 2020; Megan G. Behringer 2016; Yin et al. 2017; Qi et al. 2026). Unique structural motifs are often the cause of replication stalling and account for many of the observed genetic variants (Wei Song 2014). This is particularly true in haploid genomes that rely on non-recombinational repair (Behringer and Hall 2016). These studies used short-read, whole-genome Illumina sequencing for their analyses. Because this technology limits the window of analysis to an average of 150 base pairs, it excludes larger genetic variants from being characterized. In addition, because yeast genomes are compact and contain limited regions of heterochromatin at subtelomeric and silent mating type loci, information on rearrangements in more complex centromeres, typical of higher eukaryotes, is limited.

Recent developments in third-generation sequencing have created new avenues for detecting lower frequency structural variants (SVs) (Sereika et al. 2022; Hon et al. 2020; Poplin et al. 2018). Oxford Nanopore Technologies permit sequencing of individual chromosomal fragments as they pass through a nanopore, yielding extended reads of 10–500 kb. A second technique, PacBio Revio, sequences DNA by amplifying single molecules (typically of 10–20 kb) through multiple rounds of replication, producing a more precise consensus sequence. Standard algorithms can reliably define classes that include insertions, deletions, translocations, inversions, and duplications (Smolka et al. 2024b, 2024a). In addition, the DeepVariant AI system enables highly efficient identification of SNPs (Shafin et al. 2021). These methodologies have been used for the sequencing and structural variant analysis of multiple strains in yeast (Loegler et al. 2025; O’Donnell et al. 2023; Koen J. Dechering 1998) and have made it possible to characterize structural data from a single sequenced sample at a higher sensitivity.

*Aspergillus flavus*, a filamentous fungus, is an excellent model system for this type of analysis. *A. flavus* grows vegetatively as multinucleate mycelium (Itani et al. 2025), a feature that allows for the potential identification of rare, deleterious mutations and rearrangements that selection would otherwise tend to eliminate. *A. flavus* is also amenable to genomic analysis, and many strains have complete telomere-to-telomere sequences available (Lustig 2023; Skerker et al. 2021a). Given its relatively small, 37.5 Mb genome distributed across eight chromosomes, high-depth sequencing is readily achievable (Woods et al. 2026), increasing sensitivity for identifying rare structural variants. In addition, *A. flavus* employs both homologous and non-homologous recombination systems, typical of many eukaryotic organisms (Takahashi et al. 2012).

In *A. flavus*, centromeres and subtelomeres contain enriched A+T regions. *A. flavus* centromeres are composed of approximately 100 kb, A+T-rich region that evolved from repetitive transposable elements (Skerker et al. 2021a), bearing greater similarity to the repetitive centromeres of higher eukaryotes than to those of *Saccharomyces cerevisiae* (Bloom and Costanzo 2017). In *Neurospora* and other filamentous fungi, centromeres are embedded within heterochromatic A+T-rich sequences, shaped in part by a centromere-specific histone H3 variant (Smith et al. 2012; Smith et al. 2011). Centromere sequences on homologous chromosomes—though not on non-homologous chromosomes—generally share extensive (88–99%) homology across strains (Lustig 2023). Similarly, the subtelomere regions adjacent to the telomeric repeats span 2–20 kb and have enriched A+T base pair composition. Both regions are thought to arise through repeat-induced point mutation (RIP) (Testa et al. 2016; Cambareri et al. 1998; Lewis et al. 2009) and exist in a state of both divergence and conservation, suggesting an ongoing process that promotes and regulates recombination.

Here we look more deeply into transient structural changes across multiple cycles of growth. We previously used long-read, high-depth sequencing to identify multiple classes of low-abundance and transient inverted triplications (TRP/INVs, called iTRPs in this paper) that appear to initiate at palindromic sequences. These rearrangements were enriched in the A+T-rich centromeric and subtelomeric regions. We found that palindromes were also most prevalent in these A+T-rich centromeric regions, suggesting that their abundance—both as a source of DNA damage and as a cause of replication fork stalling—drives TRP/INV formation. These findings provide evidence that iTRPs form spontaneously during DNA replication. We postulated that such events preceded the more complex rearrangements associated with oncogenesis and other disease states in higher eukaryotes.

In eukaryotes, polydA::polydT and polydG::polydC tracts appear to play distinct mechanistic roles: polydA::polydT tracts have been associated with nucleosome exclusion and disruption of normal nucleosome positioning, whereas polydG::polydC tracts can adopt non-canonical secondary structures, such as G-quadruplexes, that are linked to replication fork stalling and altered transcription (Koen J. Dechering 1998). In humans, the main source of polydA::polydT tracts are associated with the 3’ ends of the Alu transposable element (Lustig and Petes 1984). Characterization of homopolymeric tracts in several eukaryotes has demonstrated that long poly(dA)::poly(dT) sequences are susceptible to tract-length variation, suggesting that they represent an important source of spontaneous genetic instability (Yue Zhou 2004). These tracts have historically been difficult to characterize, since their highly repetitive sequence composition remains challenging to resolve even with recent sequencing advances; nonetheless, third generation long-read technologies now make it possible to determine tract lengths with reasonable accuracy. We have employed such technologies to investigate the presence and frequency of structural variants in both homopolymeric tracts in *A. flavus*. These sequences are likely to be sites capable of DNA polymerase slippage (Gragg et al. 2002) and recombination.

Additionally, using multiple technologies, including Pepper and DeepVariant AI, we identified a transient mutation in the fifth growth cycle (G5): a single-base mutation in the putative *flbA* gene. Identification and initial characterization of this mutation highlight the value of novel bioinformatic technologies in detecting sequence changes that are difficult to capture through wet-lab genetic testing alone.

Our results support four general categories: (a) SNPs; (b) precise deletions associated with flanking regions of homology; (c) clusters of short deletions heavily enriched within high-A+T centromeric and subtelomeric regions; and (d) deletions and expansions within a subset of homopolymeric tracts, reproducible across independent datasets. We identified 70 long (> 44 bp) homopolymeric tracts dispersed throughout the genome in noncoding regions, a feature common among the *A. flavus* species we surveyed within the phylum Ascomycota. These findings demonstrate that deletions and expansions, particularly within regions of simple-sequence DNA, constitute a significant class of spontaneous variation in *A. flavus*.

## RESULTS

### Development of an *Aspergillus flavus* model for the Detection of Structural Variants During Vegetative Growth

We endeavored to develop an *A. flavus* model system to probe spontaneous genetic changes during vegetative growth in the absence of overt selection. This system has the advantage of identifying potentially deleterious mutations in multinucleate cells. To this end, we cultured *A. flavus* for 15 cycles of mycelial growth (each cycle consisting of a week of growth), followed by dilution and continued vegetative growth.

Libraries of DNA isolated from mycelial cells after 0, 5, 10, and 15 cycles of growth (G0, G5, G10, and G15) were constructed and subject to either nanopore (G0-G15) or PacBio Revio (G15) sequencing, and assemblies from each of these DNAs were constructed. We first wanted to test whether the available algorithms for analyzing sequencing data were sufficiently sensitive to identify the genetic basis of a phenotypic change in mycelial morphology (from yellow/green to white mycelia and conidial development) between the fifth (G5) and tenth (G10) weeks of culturing (Figure 1A). We hypothesized that the basis of this morphology was a change in standard conidial structure and density. We therefore measured the density of conidia directly in a hemocytometer, using brightfield microscopy for the original NRRL 3357 G0 strain and for the NRRL 3357 strain after 15 cycles (G15). Cells were plated in both light and dark conditions, since light can regulate asexual spore formation in filamentous fungi through the regulation of the *brlA*, *fluG*, and *flbA*-*E* genes in *Aspergillus nidulans*, when plated on normal rich YGT (Yeast Extract, Glucose, Trace Elements), PDA (Potato Dextrose Agar) and WKHM (Wickerham) media (Ruger-Herreros et al. 2011; Son et al. 2023). We found that G0 conidia were present at high levels (300,000-500,000 conidia/mm2) in both light and dark conditions, but that G15 had only residual levels of conidia. Interestingly, this was media-dependent, since when plated on glucose minimal media (GMM), cells showed phenotypic changes in morphology, but only in the presence of light (**Figure 2**).

**Fig 1:**
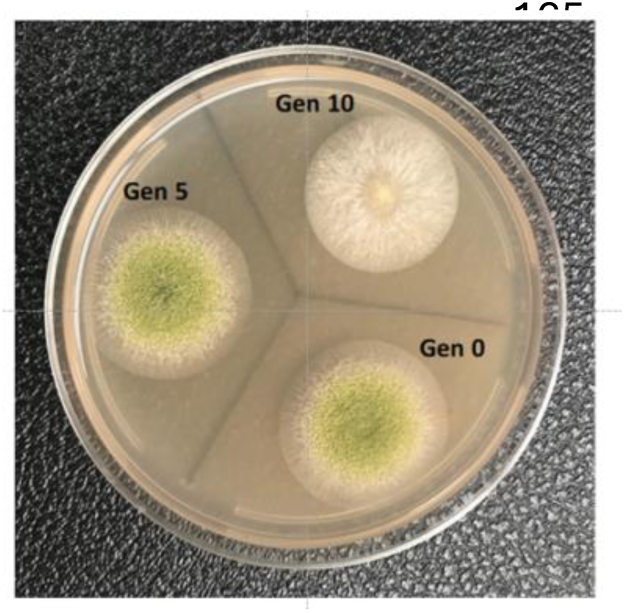
A Phenotypic Alteration Arising between the 5^th^ and 10^th^ Subculturing is Associated with Reduction in Conidial Asexual Spore Formation. Morphology of G0 (Gen 0), G5 (Gen 5) and G10 (Gen 10)

**Fig 2:**
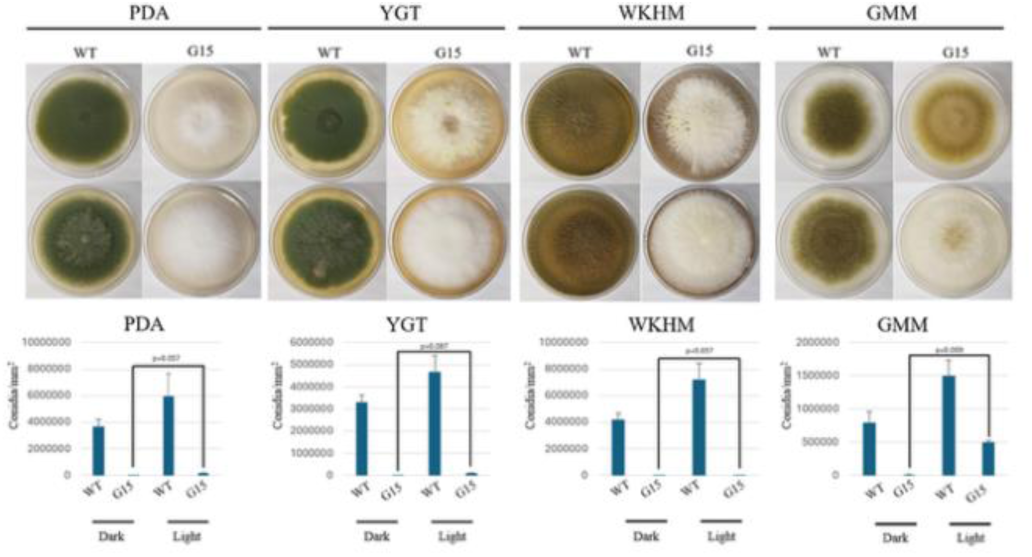
Quantification of asexual spore production. (A) Photographs of the NRRL3357 wild type and G15 mutant strain after growth on PDA, YGT, Wickerham (WKHM), and GMM media for 7 days under light and dark conditions at 30°C. (B) Quantification of conidiospores in panel A. Error bars represent standard error and letters over the bars indicate statistical significance. Statistical significance (p-values; One-Way ANOVA-Games-Howell post hoc test) between the dark and light conditions for the G15 mutant are shown for each media type. The difference is highly significant under GMM media conditions.

Hence, both nutrient conditions and the activation of specific gene products may be involved in this selective response (Figure 2, Supplementary Figure 1). Our initial analysis using standard DNA sequencing algorithms (Sniffles 2) to detect nuclear and mitochondrial structural variants found no differences in deletions, insertions, inversions, or translocations between G0, G10, and G15 cultures (data not shown).

To detect single-nucleotide polymorphisms (SNPs) that may have arisen during growth, we compared the highest-quality PacBio reads to each ONT-derived assembly using the Pepper/Deep Variant AI system (Shafin et al. 2021). PacBio was used for these analyses since reads were more closely identified (99.9%) to the assembly sequence. The Pepper data were filtered to examine only SNPs, and the results were annotated with potential coding regions. Only one mutation was identified in a coding region when comparing G15 PacBio to the G0 ONT assembly: a G-to-A mutation in a the AFLA 134030 putative protein that shifts Asp to Asn at amino acid 672 (Figure 3). To examine the presence of the mutation in G5 and G10 cells, we compared the G15 PacBio to each respective G5 and G10 ONT-derived assembly. While the wild-type allele was present in G5 cells, it was no longer detectable in the G10 assembly, consistent with the appearance of the mutant.

**Fig 3.**
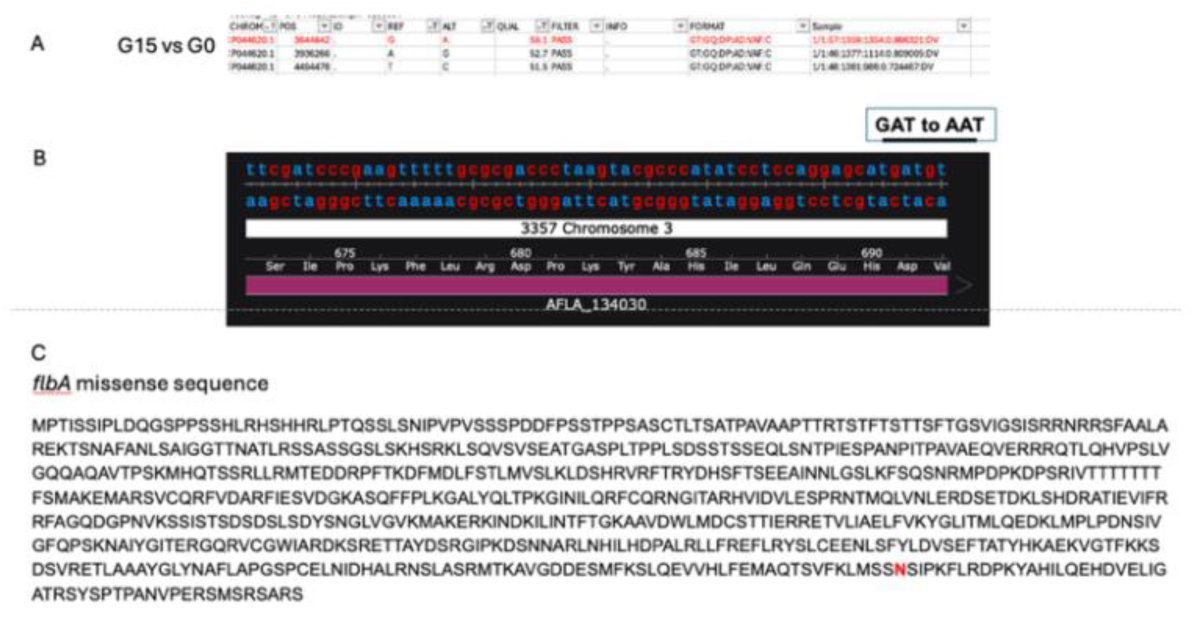
G15 and G10 Cells Contain a Missense Mutation in AFLA_143030 Not Present in G0 and G5 Cells (A). The G15 PacBio data set was compared to G0, G5, and G10 assemblies using Deep Variant-Pepper, and the results were compared for single-nucleotide polymorphisms with a variant allele frequency of over 90%. Only one sample had a variant allele when comparing G15 reads to G0 and G5 assemblies, but not to a G10 assembly in the vast majority of cells (shown in red). This mutation mapped to a G to A transition in the putative gene product AFLA_134030 on chromosome 3 (B).

### Further BLASTP, Conserved Domain Database (CDD), and AlphaFold3 analyses reveal that *AFLA_134030* is a structural homolog of *A. nidulans flbA*

Although not identified yet in *A. flavus,* the *flbA* gene, as noted above, is a regulator of conidial formation and asexual spore formation in *A. nidulans* (Ruger-Herreros et al. 2011). Based on the light-activated phenotype previously described in *A. nidulans*, we tested whether this AFLA_134030 is a missense mutation in the *flbA* gene. We employed three bioinformatic tools to investigate the identification and impacts of the missense mutation. Upon identifying the specific missense mutation in *AFLA_134030*, we used NCBI’s BLAST P function to determine the amino acid homology between the *AFLA_134030* and the *A. nidulans flbA* gene (Altschul et al. 1997). We found 82% homology between the two sequences, with the *A. flavus* gene being 9 amino acids shorter than that of *A. nidulans*, (719 and 728 bp respectively) (Supplemental Figure 2). Although this indicated high homology between the two proteins, we endeavored to investigate similarities at the structural level.

Using the CD database, we identified strong conservation between the C-terminal amino acids of *A. flavus flbA* gene and AFLA_134030 amino acids 545-683bp (Supplemental Figure 3) (Marchler-Bauer and Bryant 2004), within the *A. nidulans flbA* G protein signaling domain. The missense mutation at amino acid 672 falls within this domain.

To examine the structural homology further, we uploaded the *A. flavus* and *A. nidulans* target sequences into AlphaFold 3 (Abramson et al. 2024a, 2024b), and compared structural similarity using the Chimera X Matchmaker structural program. This analysis indicates that over the five most probable predicted structures for proteins, the structure of the two proteins was similar, with a Root Mean Square Deviation (RMSD) of .467 Angstrom difference by the Needleman-Wunsch algorithm (Supplemental Figure 4). Given that RMSDs of 1-2 angstroms are considered an acceptable margin of error between similar proteins, this finding highlights the probability that the putative *A. flavus* FlbA (pAf FlbA) is a structural homolog of *A. nidulans* FlbA.

This study demonstrates the utility of AI-assisted SNP analysis coupled with by a combination of phenotypic and bioinformatic approaches for detecting mutations in real time growth through the identification of an A. flavus structural and functional homology to A. nidulans FlbA.

### The Largest Class of Precise Deletions is Flanked by Short Repeats and is Found in Both Coding and Noncoding Regions During Vegetative Culturing

Having previously identified inverted triplications (iTRPs) that arise from palindrome-associated replication stress by ONT sequencing, we applied long-read, high-depth sequencing to determine additional classes of spontaneous structural variants, such as deletions and expansions (Woods et al. 2026). To identify short deletions or expansions of ≥ 18 bp, we used PacBio Revio sequencing technology.

We found that PacBio reads had a very low overall error rate (0.1%), with even a relatively low rate (∼1%) in highly AT-rich sequences (Supplementary Figure 5). ONT sequencing, on the other hand, while valuable for identifying large-scale rearrangements, contained too many random misreads shorter than 45 bp and was therefore not sufficiently sensitive to detect events verified in a small number of sequencing reads (data not shown).

The class with the highest read frequency (copy number), ranging from 5 to 37, is precise deletions (i.e., deletions in which reads could be verified to arise from specific junctions), with deletion sizes varying widely from 18 to 15,136 bp. Five of these deletions occurred in putative genes, while two were present in noncoding regions (Table 1). Algorithmic and Integrative Genome Viewer (IGV) analyses yielded somewhat different copy numbers, in part due to the exclusion of short reads from the IGV and to errors in the algorithmic output.

**Table 1.** Precise Deletions Detected by PacBio Sequencing. ALG, algorithmic, IGV, Integrative Genomic Viewer. Red, closely lying duplicated copies of designated genes * Genes include AFLA_009201, 009202, 009203, 009204, 009205, 009206, 009207 and F9C07_2108266. The chromosome 7 large insertion is flanked by the sequence CTGTTGTATG, present starting at position 2916251 and position 292.

| CHR | COORDINATES/<br>LENGTH (BP) | COPY<br>NUMBER<br>(ALG) | COPY<br>NUMBER<br>(IGV) | SHARED<br>JUNCTIONS | VARIANT<br>JUNCTIONS | GENE<br>REGION | JUNCTIONAL<br>HOMOLOGY |
| --- | --- | --- | --- | --- | --- | --- | --- |
| 1 | 3568043-3568060 | 11 | 8 | 7 | 1 | NC | NO |
|  | (18) |  |  |  |  |  |  |
| 4 | 1829096-18299605<br>(510) | 10 | 10 | 8 | 2 | AFLA_0<br>02414 | YES |
| 4 | 3404527-3404563<br>(34) | 5 | 5 | 5 | 0 | AFLA_0<br>11877 | YES |
| 4 | 3404936-3404971<br>(34) | 5 | 5 | 5 | 0 | AFLA_0<br>11877 | YES |
| 4 | 4614492-4614590<br>(99) | 20 | 14 | 14 | 0 | AFLA_0<br>11438 | YES |
| 5 | 289727-289825<br>(99) | 45 | 37 | 37 | 0 | FYC07_<br>2284313 | YES |
| 5 | 2712422-2727584<br>(15136) | 18 | 9 | 9 | 0 | Multiple<br>* | NO |
| 7 | 2916263-2927859<br>(11597) | 20 | 17 | 15 | 2 (3) | NC | YES |

In most cases, the deleted regions were flanked by short repeated sequences (Figure 4), and in one case, the deletion was itself associated with a duplication (Figure 4C). These data suggest that such flanking repeats are associated with deletions and that recombination between such short homologies may be involved in the excision of the deleted sequences during vegetative growth. In the 11.6 kb deletion, we also observed a short flanking repeat. In the 15.1 kb deletion, the left breakpoint was within a predicted intron, and the right junction was in a noncoding region that eliminated eight genes.

**Fig 4.**
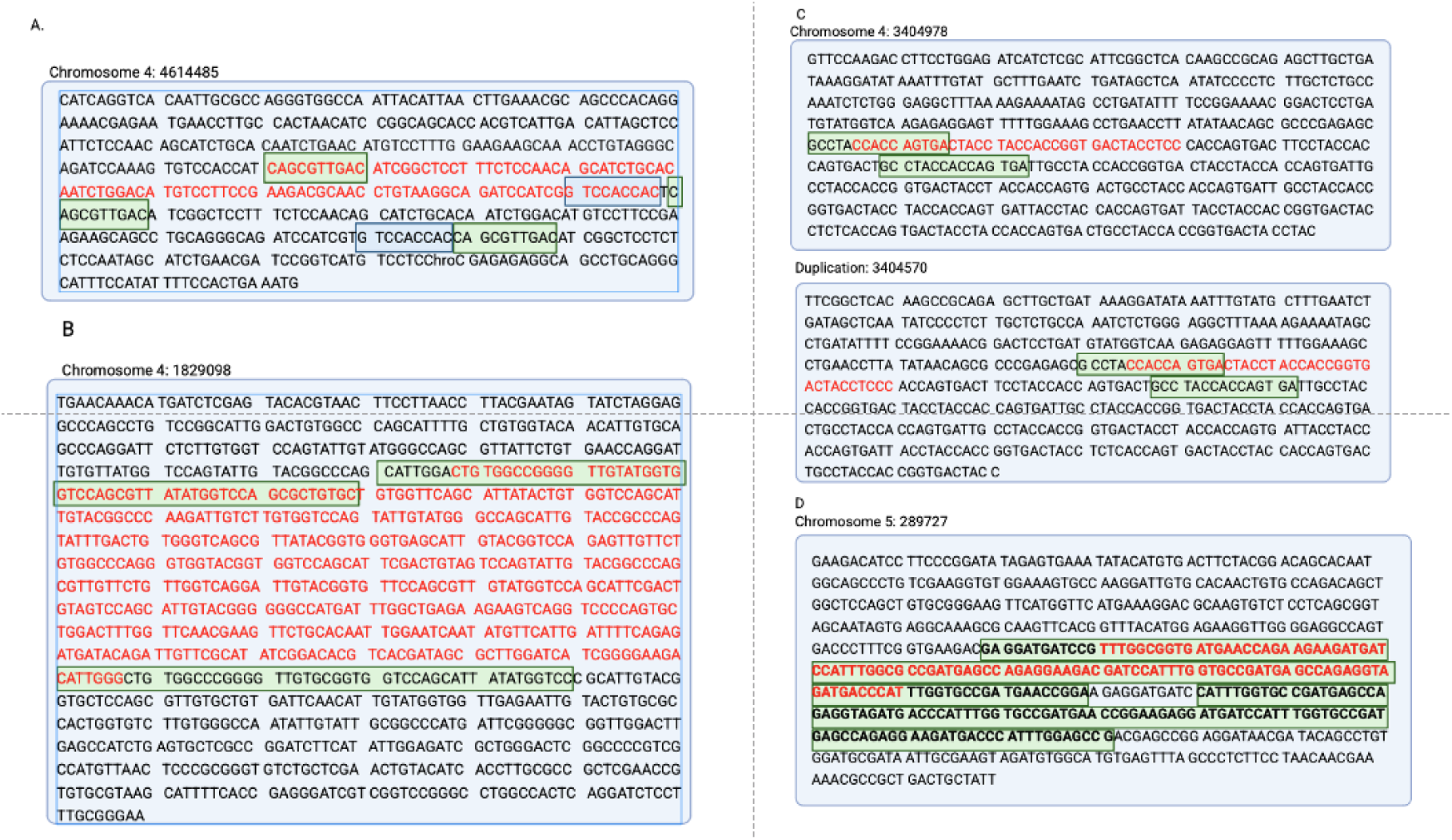
The Most Abundant Class of Precise Deletions Is Correlated with Short Repeats Flanking and Within the Deleted Sequence. A BLAST analysis of the sequences surrounding the deletions to themselves revealed short repeats. A subset of the results is shown in boxes A-D with a green or blue shaded color showing each set of short repeats. Other repeats were also found within this region (data not shown)—the rearrangement in C results in a gene duplication accompanied by deleted sequences at homologous sites.

IGV analysis revealed that, in three cases, the deletion endpoints showed microheterogeneity, suggesting that they likely occurred in multiple independent events. A careful examination of putative structures and repeat patterns revealed no predicted characteristics of folding as determined by UNAfold algorithm (Markham and Zuker 2008), except for the paired 34 bp deletion (AFLA_011877) (data not shown).

In summary, these data suggest that the presence of short repeated sequences has a direct or indirect effect on the generation of spontaneous deletions during vegetative growth (see Discussion).

### Clustered Short Deletions Are Enriched in AT-rich Subtelomeric and Centromeric Sequences

We have previously shown that the clustering of palindromes in centromeric and subtelomeric regions helps to drive the formation of inverted triplications (Woods et al. 2026). Our earlier studies also revealed that direct and inverted repeats are also enriched in these regions. Since such repeats are prone to genetic instability, we next sought to identify regions susceptible to the formation of indels, in particular short deletions. Several additional steps were included to minimize the misidentification of short deletions, particularly in regions of simple-sequence DNA, where false algorithmic calls or poorer DNA sequencing quality can lead to misinterpretation. First, a Sniffles 2 algorithm, empirically optimized to reduce errors in simple sequence regions, was employed as described (Smolka et al. 2024b, 2024a). Second, an IGV visual analysis was conducted to eliminate false positives containing multiple linked events (Thorvaldsdottir et al. 2013). By these means, we identified two additional broad classes: a) clustered short deletions, and b) expansions and contractions in homopolymer tracts. We found that the Sniffles 2 algorithms often identified clustered sequences as the sole change in a short DNA region from one growth cycle to the next. We eliminated low-quality DNA regions as described in Materials and Methods.

The deletions identified were separated into three categories based on their genomic site; the ∼100 kb A+T-rich centromeric domains, the AT-rich elements (ATEs) located predominantly within the 100 kb subtelomeric regions (and in several cases in non-subtelomeric locales), and the non-ATE regions dispersed throughout the genome (Table 2). Thirty clusters (2-4) of short deletions, ranging in size from 16 bp to 48 bp, were present in centromeres, with an average deletion size of 23 bp. In this region, the average AT-content was 90%, while the deleted sequences ranged from 90 to 97%, with an average AT content of 95% (Table 2A, Supplemental Table 1A). Thirteen additional clusters were found in subtelomeric (and other) regions that had a similar higher-than-average A+T composition (Table 2B, Supplemental Table 1B). In contrast, only 28 other clusters were observed in the remainder of the genome with average deletion sizes of 32 bp. (Table 2C, Supplemental Table 1C). We estimate that deletions in centromeric and ATE regions are 34- and 48-fold more frequent than in dispersed loci, respectively (Table 3). Consistent with their high AT-content, UNA-fold analysis of the region surrounding the deletion sites reveals highly paired structures (data not shown).

**Table 2.**
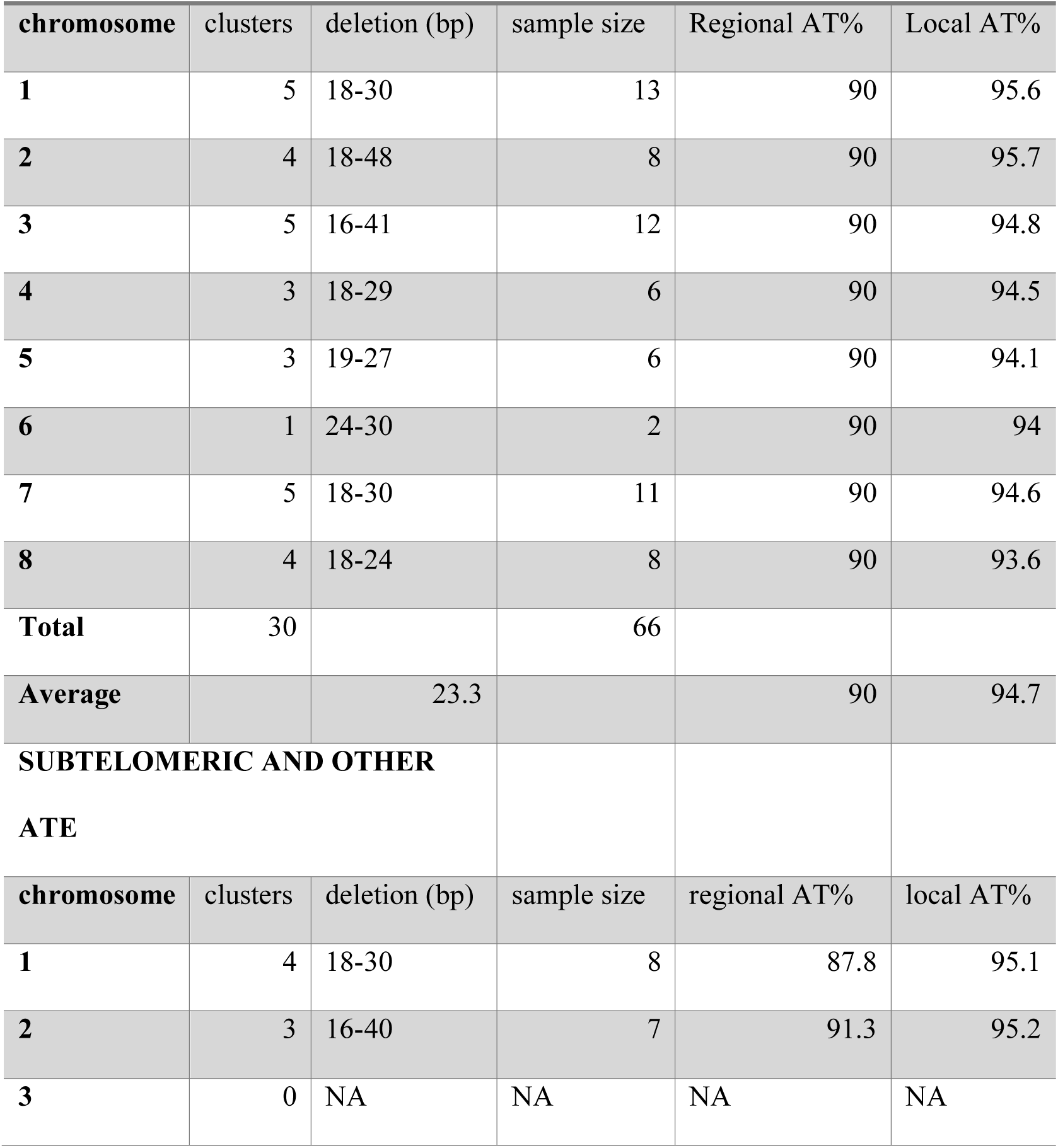
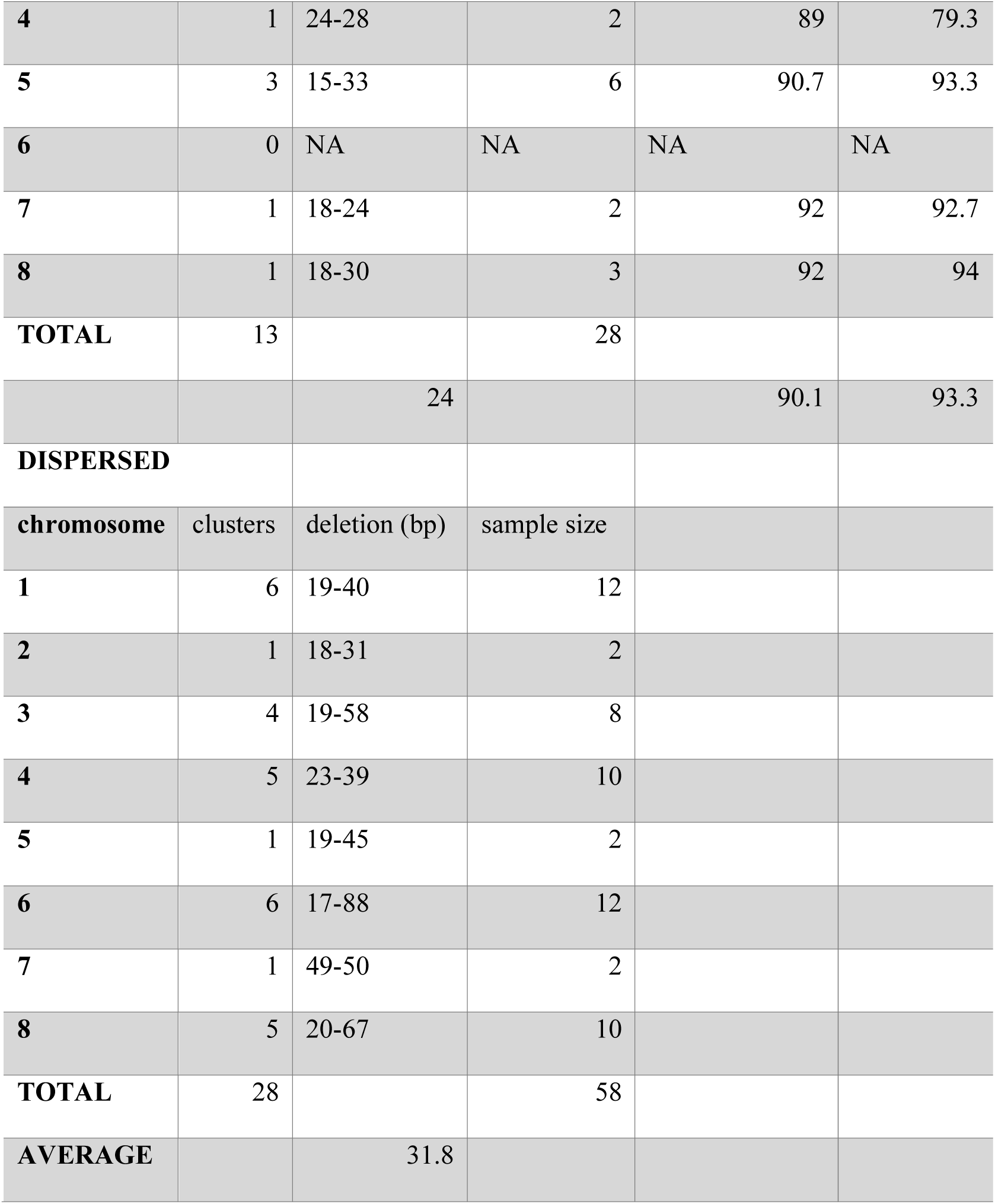
Aggregate Results of Clustered Deletions. Summary of data shown in Supplementary Tables 1A-1C describing the results from (top) centromeric, (middle) subtelomeric and other ATEs, and (bottom) dispersed domains. The results are presented for each of the chromosomes, and show the number of clusters, the range of deletion sizes and the aggregate read numbers. For centromeric and subtelomeric regions, the AT content of the entire centromere or ATE is given as the regional %AT and the 200 bp region that includes the deletion is given as the local %AT.

CENTROMERE
| chromosome | clusters | deletion (bp) | sample size | Regional AT% | Local AT% |
| --- | --- | --- | --- | --- | --- |
| 1 | 5 | 18-30 | 13 | 90 | 95.6 |
| 2 | 4 | 18-48 | 8 | 90 | 95.7 |
| 3 | 5 | 16-41 | 12 | 90 | 94.8 |
| 4 | 3 | 18-29 | 6 | 90 | 94.5 |
| 5 | 3 | 19-27 | 6 | 90 | 94.1 |
| 6 | 1 | 24-30 | 2 | 90 | 94 |
| 7 | 5 | 18-30 | 11 | 90 | 94.6 |
| 8 | 4 | 18-24 | 8 | 90 | 93.6 |
| **Total** | 30 |  | 66 |  |  |
| **Average** |  | 23.3 |  | 90 | 94.7 |

**Table 2.** Aggregate Results of Clustered Deletions.
| chromosome | clusters | deletion (bp) | sample size | regional AT% | local AT% |
| --- | --- | --- | --- | --- | --- |
| 1 | 4 | 18-30 | 8 | 87.8 | 95.1 |
| 2 | 3 | 16-40 | 7 | 91.3 | 95.2 |
| 3 | 0 | NA | NA | NA | NA |
| 4 | 1 | 24-28 | 2 | 89 | 79.3 |
| 5 | 3 | 15-33 | 6 | 90.7 | 93.3 |
| 6 | 0 | NA | NA | NA | NA |
| 7 | 1 | 18-24 | 2 | 92 | 92.7 |
| 8 | 1 | 18-30 | 3 | 92 | 94 |
| **TOTAL** | 13 |  | 28 |  |  |
|  |  | 24 |  | 90.1 | 93.3 |
| **DISPERSED** |  |  |  |  |  |
| **chromosome** | **clusters** | **deletion (bp)** | **sample size** |  |  |
| 1 | 6 | 19-40 | 12 |  |  |
| 2 | 1 | 18-31 | 2 |  |  |
| 3 | 4 | 19-58 | 8 |  |  |
| 4 | 5 | 23-39 | 10 |  |  |
| 5 | 1 | 19-45 | 2 |  |  |
| 6 | 6 | 17-88 | 12 |  |  |
| 7 | 1 | 49-50 | 2 |  |  |
| 8 | 5 | 20-67 | 10 |  |  |
| **TOTAL** | 28 |  | 58 |  |  |
| **AVERAGE** |  | 31.8 |  |  |  |

**Table 3.**
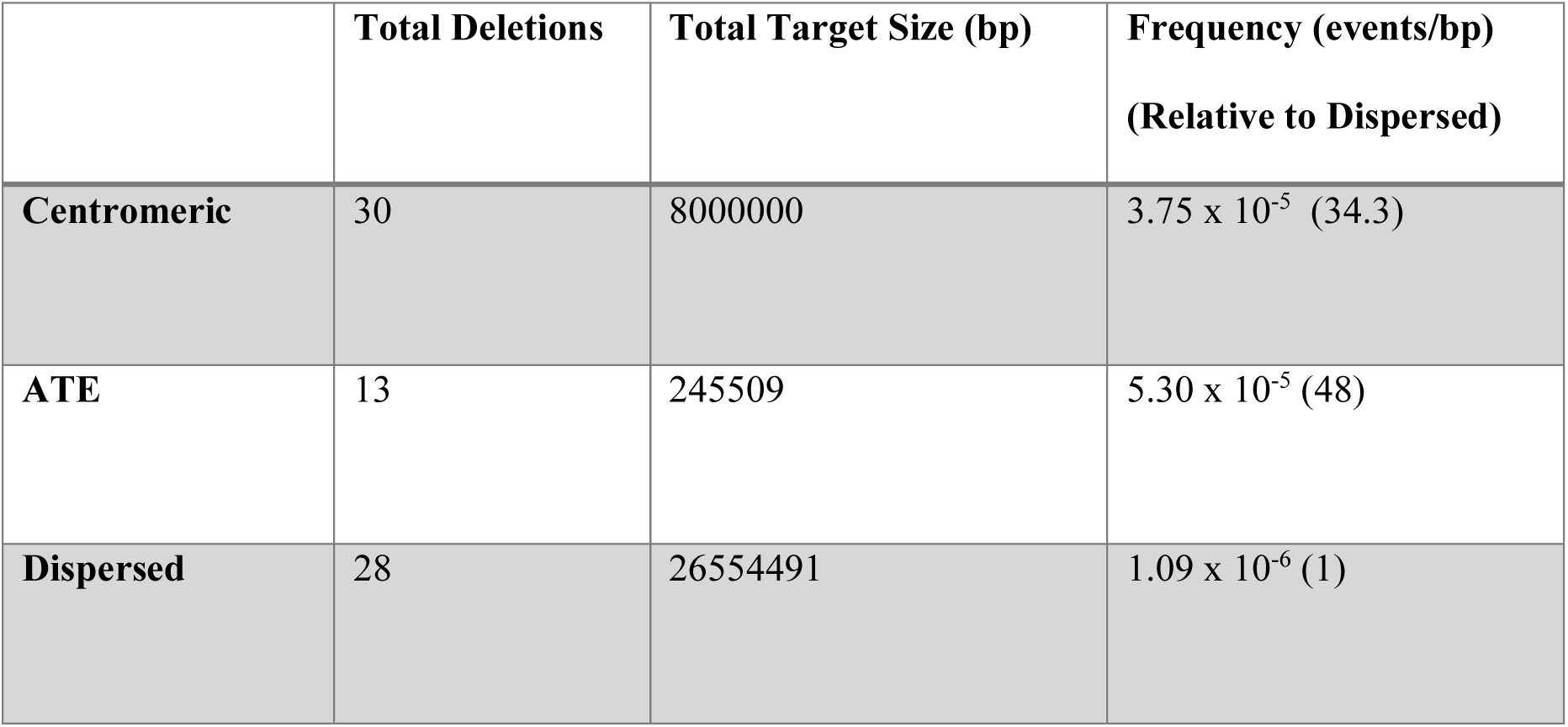
Frequency of Deletion Clusters in Centromeric, ATE, and Dispersed Domains. Shown here are the total number of deletions in each domain, the aggregate size of the assayed domain and the deletion frequency.

### CENTROMERE

#### PacBio Sequencing Reveals Deletions within Homopolymer PolydA::PolydT and PolydG::PolydC Tracts

The length and stability of homopolymeric tracts have been refractory to analysis because ONT and previous PacBio sequencing technologies were not sufficiently accurate to analyze them. We have, however, found that the PacBio Revio sequencing methods can determine their length. The IGV scans reveal very few changes beyond a systematic assembly error in polydG::polydC tracts. The variation between reads is slight compared with previous studies, approximately 5 bp (data not shown).

Based on this stability, we have surveyed the landscape of homopolymer tracts throughout the *A. flavus* genome that are 20 bp or larger. 213 polydA::polydT tracts, ranging in size from 20-75 bp, and were dispersed among both introns and noncoding regions of every chromosome, with the exception of chromosome 7. 53 polydG::polydC tracts ranging from 18 bp to 52 bp are also present on all chromosomes, except for chromosome 8 (Figure 5 A, B; Supplementary Table 2). A genomic map of the tract distribution on the chromosomes is shown in Figure 6.

**Fig 5.**
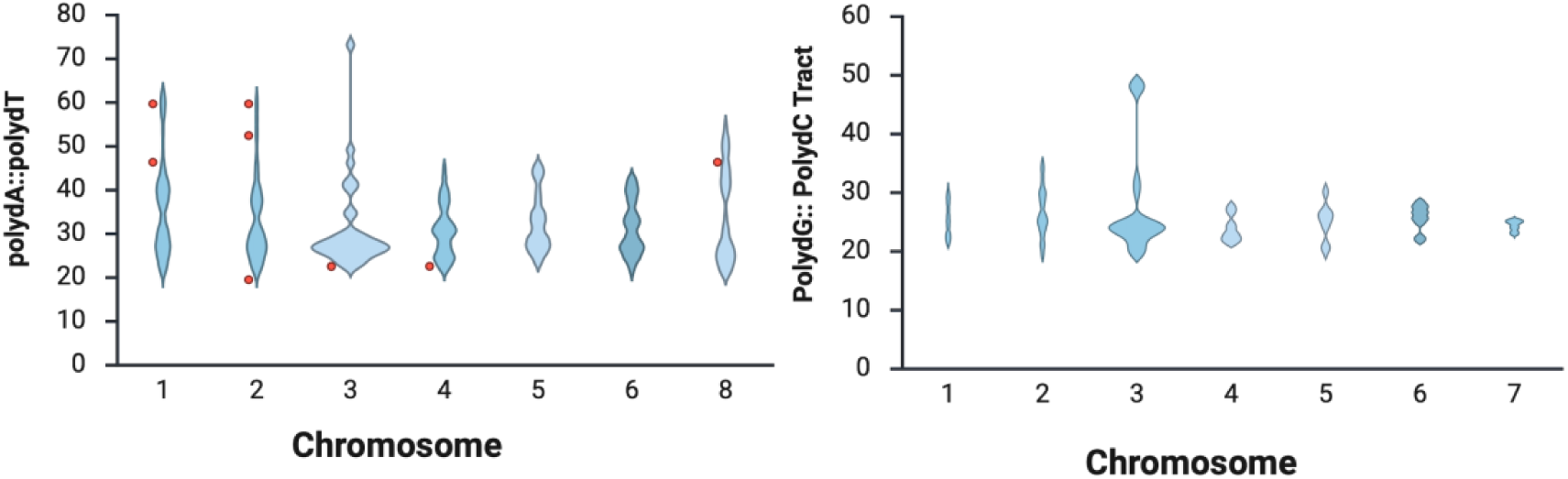
The Distribution of Homopolymer Tract Sizes Across A. flavus Genome (From Supplementary Table 2). Violin plots of the length (y-axis in bp) of (left) the polydA::polydT or (right) polydG::polydC homopolymers on each chromosome (x-axis). The sizes of the tract that were subject to deletion in these studies are shown by red dots. Homopolymeric polydG::polydC lengths may be slightly shifted due to a common occurrence of assembly errors after DNA sequencing.

**Figure 6.**
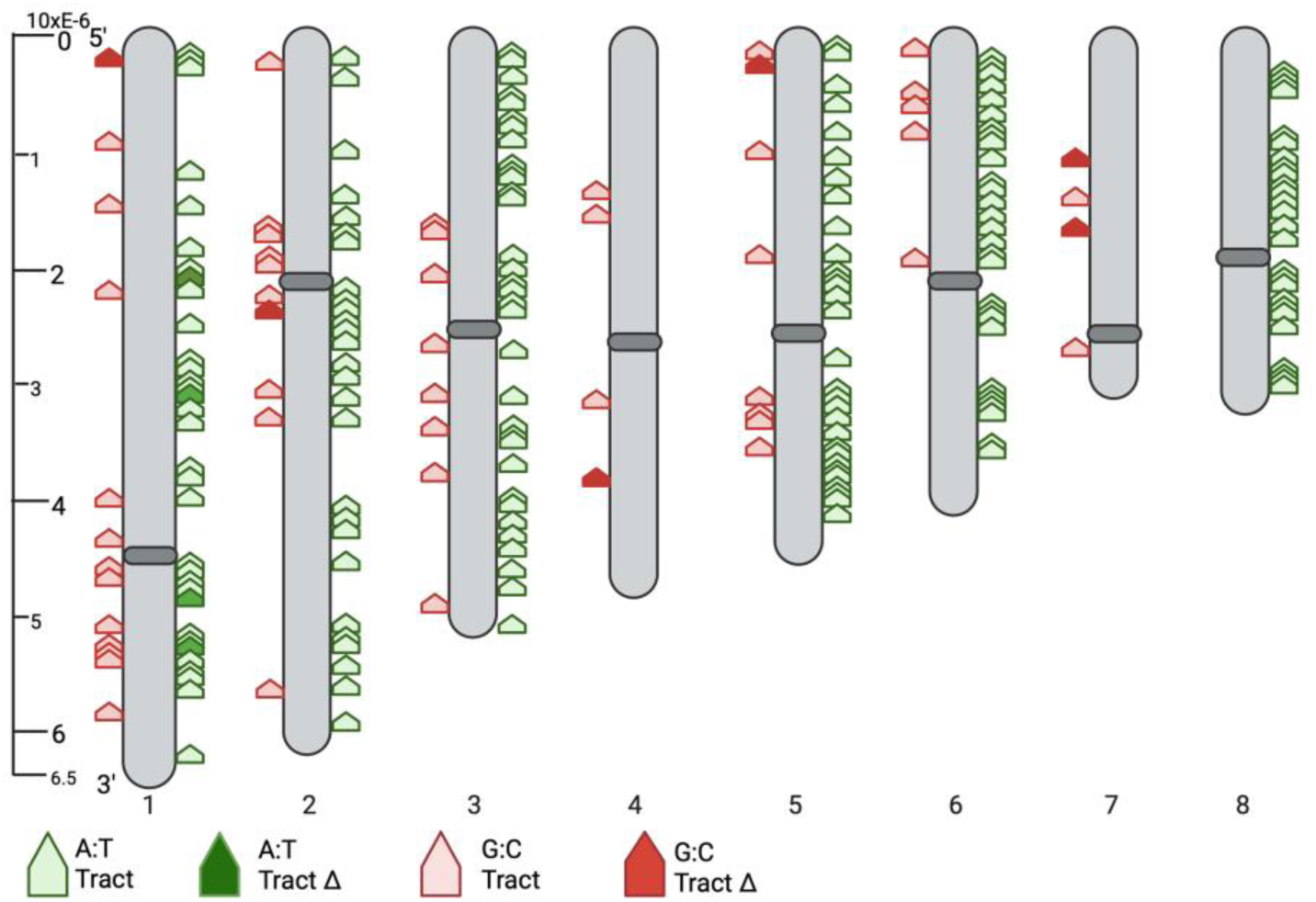
A Genomic Distribution Map of PolydG::PolydC (GC) and PolydA::PolydT (AT) Sequences and Deletion Sites. The deletion sites are shown by the filled red or green pentagons.

To determine the generality of this finding across evolutionarily related filamentous fungi, we surveyed the genomes of different species from published datasets that were sequenced using high-quality PacBio HiFi and ONT methodologies, as judged by the full sequence of the A+T-rich centromere. Interestingly, we found a distribution that contained longer tracts (≥ 44 bp) in all ascomycete’s filamentous fungi, although differing in abundance (Figure 7, Supplementary Tables 3 (Source Data) and 4). These polydA::polydT homopolymeric tracts were unassociated with assayable signatures of transposable elements using the Censor algorithm, suggesting that these tracts are not linked to the retrotransposition of polyadenylated RNAs (data not shown). The biological functions of these homopolymer tracts are unknown, but both are likely to be mediated by the secondary functions of non-canonical DNA structures (see Discussion).

**Figure 7.**
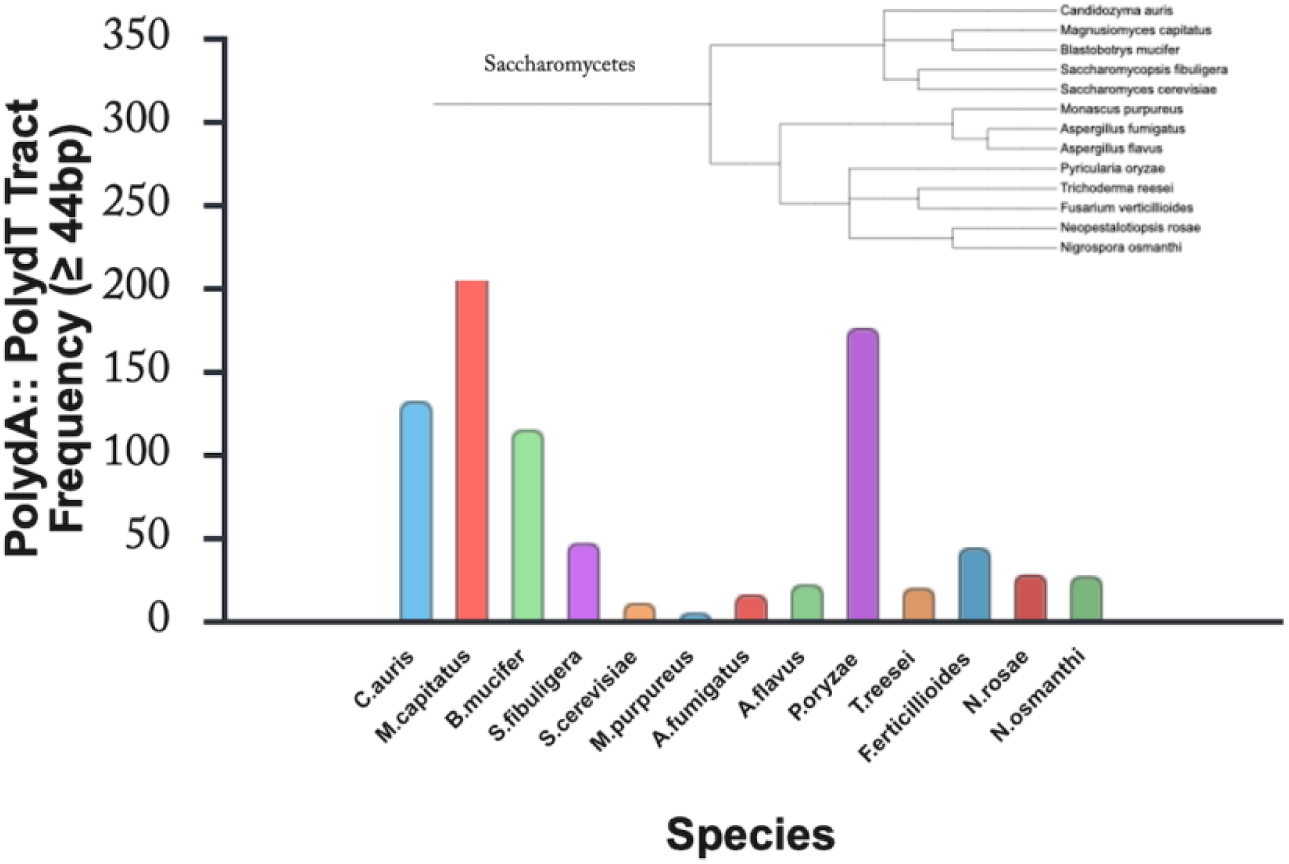
Long PolydA::PolydT Tracts Are Common Among the Saccharomyces Species Related to A. flavus. Datasets of high-quality sequences mined from the NCBI database were assayed for long polydA::polydT sequences listed, for convenience, as sizes greater than or equal to 44 bp. The number of tracts is shown on the y-axis and the species tested on the x-axis. The insert shows the phylogenetic similarity of the strains employed.

Given the ability to reliably measure the length of homopolymer tracts, we sought to determine their stability by identifying structural variants using the same steps employed for clustered deletions. We found that 9 of the 213 polydA::polydT homopolymer tracts contained deletions corroborated in 2-123 independent reads from G15 cells. The length of the deletion was estimated in a subset of BLAST analyses, but in some cases varied from the algorithmically determined estimate (Supplementary Figure 6A). However, since we found a population with slightly different deletion sizes, there may be a small sequencing or analysis error contributing to this variation. Alternatively, multiple events may occur within these tracts, coupling a large deletion with more minor changes. Further, to rule out the possibility that these deletions were generated during DNA sequencing, we compared the results with a second independent sequencing of G15 cells with reduced read density. We found that six of the nine reads in the second G15 dataset were also deleted, and they were present at the same relative frequency despite the 50% decrease in depth (Table 4A). These data indicate that many, if not all, of these deletions were due to a preexisting defect in a subpopulation of the DNA. Deletions were not exclusively associated with the longest tracts, as indicated by the red ticks in the violin plot of Figure 2A.

**Table 4.**
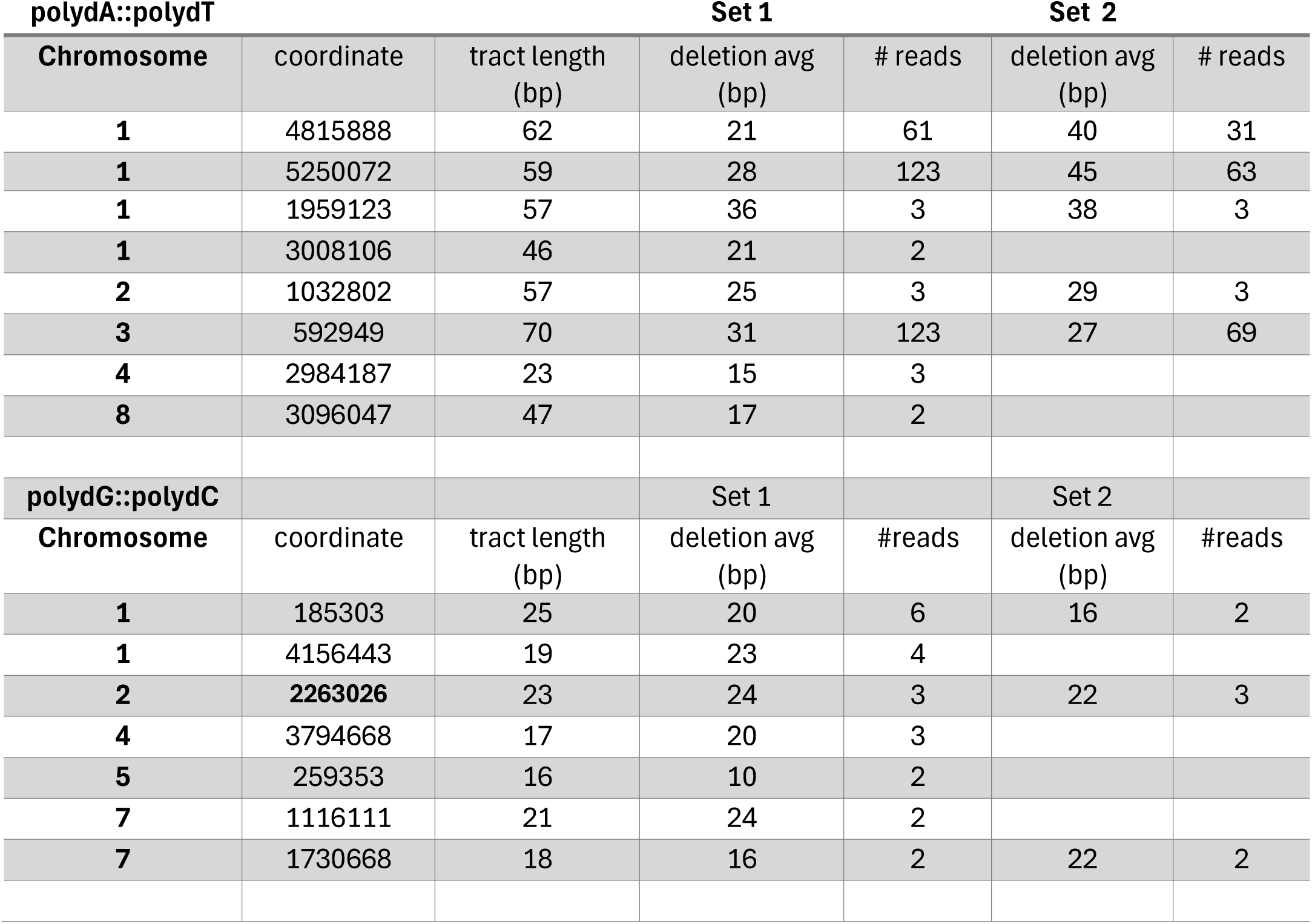
G15 Deletions in Homopolymeric Tracts. The characteristics of homopolymeric deletions shown here are derived from two G15 sequence data sets, with set 2 having approximately 50% of the coverage of the first dataset used for the rest of this study. The bold font indicates the polydG::polydC tract identified also by nanopore sequencings (data not shown).

A similar analysis was carried out for polydG::polydC tracts. We identified that 7 of the 53 polydG::polydC homopolymer tracts contained deletions that could be corroborated from 2-6 reads from G15 cells. The lengths of the tracts in this case were corrected for a small assembly error exhibited in all reads. The length of the deletion was verified by using a BLAST analysis (Supplementary Figure 5B). In this case, only three deletions were corroborated by the second independent database, probably because of the lower read depth observed for the polydG::polydC candidate deletions, and no additional deletion sites were found in the second set (Table 4B), again arguing for a preexisting defect in a subpopulation of the DNA.

We wanted to know whether the identified deletion sites could also be prone to expansion. We found two polydA::polydT expansions of 10 bp at Chr1/4815888 and Chr1/5250072, and two 8 bp expansions in Chr3/592949. Similarly, we found two polydG::polydC expansions at Chr1/ 4156443 (24-29 bp), Chr2/ 2263026 (22-23 bp), and Chr7/ 1730668 (25-35 bp) (data not shown). These data may represent expansion events mediated by a similar process (e.g., DNA polymerase slippage) or by a unique process relative to deletions. While expansions are infrequent relative to deletions, they further indicate the instability of these regions. We did not, however, explore expansion at the genomic level.

In summary, homopolymer tracts can now be assessed using PacBio Revio sequencing, and this assay can be useful both for determining their distribution in subsequent studies of their potential functions and for assessing their genetic instability.

## DISCUSSION

In this study, we explore the landscape of genetic variation in the model system *Aspergillus flavus* in the absence of an overt selection. This type of analysis is important for defining the types of genetic instabilities that arise and can subsequently form more complex rearrangements under selective conditions. In addition to the previously reported inverted triplications, we identified several additional sources of structural variants as deletions and, in some cases, expansions. This analysis used PacBio sequencing, which had 99.9% read identity to the assembly. We did not observe any translocations or inversions.

First, we found a class of unique 18 bp-15 kb deletions substantiated by multiple reads (defined algorithmically as precise deletions) in both genic and non-genic regions of the genome. Most of these classes were associated with numerous short direct repeats, including those flanking the deletion region. These range in size from very short 10 bp sequences to larger 110 bp sequences. While these results are descriptive, they suggest that these direct repeats may be involved in deletion, either through simple excision or deletion through microhomology-mediated end joining (Meyer et al. 2015). The finding that one of these deleted regions on chromosome 4 was also associated with a partial gene duplication may argue for a general instability of these regions.

Second, we observed a class of deletions that we have referred to as clustered deletions. This class contains short deletions defined by multiple reads spanning a 200 bp segment of otherwise high-quality DNA. These are not derived from a single genetic change, but instead suggest an increased probability of deletions in specific regions. Such events were found in both centromeric, subtelomeric (and other) ATEs and in dispersed regions. Interestingly, we found that deletions in centromeric and subtelomeric regions occurred in areas with extremely high AT content. In centromeric regions with an average AT composition of 90%, deleted regions were, on average, 95%. At this AT-concentration, the probability of forming extended paired regions, as determined by UNAfold, is very high We speculate that such long palindromes are subject to DNA repair during DNA replication. We note that the quantitative (34-fold) increase in deletions in centromeres is similar to the 35-fold increase observed in inverted triplications, possibly due to a similar underlying process driven in part by the enrichment of palindromes in centromeric regions (13). These data raise the possibility that the repair of palindromes into inverted triplications and deletions may be both the consequence of palindrome instability, in the latter case processed by exonucleolytic cleavage coupled with either non-homologous or microhomology-mediated end joining (Chang et al. 2017).

Third, we have identified both deletions and expansions in homopolymeric tracts. Homopolymeric tracts are present throughout eukaryotes but have been technically refractory to third-generation sequencing and genomic analysis. These results, using the most recent PacBio technology, constitute the first genomic map of simple homopolymeric tracts, which may subsequently be used as a reference for their association with transcriptional and recombinational regulation. Past analyses of homopolymeric tracts have indicated their association with the reverse transcription of polyadenylated RNA during retrotransposition (for example, those associated with Alu repeats). However, we have found no linkage between transposable elements and footprints in *A. flavus*. Even though polydA tracts arise ancestrally from transposable elements, their retention suggests an additional function acquired during evolution. Both polydA and polydG tracts are interspersed throughout the genome. Both polydA and polydG tracts can be found in noncoding regions, upstream and downstream of putative genes, as well as in introns.

PolydA::polydT tracts have been associated with nucleosome disruption through the formation of non-canonical DNA structures that may render them susceptible to numerous enzymatic systems and recombination. Longer tracts may also lead to reorganization of nucleosome positioning and phasing. The commonality of these tracts in filamentous fungi may reflect a unique function, such as the regulation of secondary submetabolite gene clusters. PolydG::polydC tracts are found in promoter regions where they are also not packed efficiently into nucleosomes and can also fold into G-quadraplex structures (and sometimes triplex DNA) that can play a role in transcription and replication stalling through monovalent-stimulated Hoogsteen G-G base pairing. Such tracts similarly act as hotspots for genetic instability and mitotic recombination, which can further be influenced by selective methylation and demethylation (Campbell et al. 2025). The description of these tracts across the genome will allow a study of the effects of individual tract deletions on the expression of upstream or downstream genes and gene clusters, as well as on their roles as sites of genetic instability.

Sequencing analysis has identified both deletions and expansions of the polydA::polydT homopolymers. These were substantiated by two independent sequencing runs and were verified by many reads. At this time, it is difficult to determine whether these were multiple independent events or the expansion of a single event. The sites for deletions were also capable of small expansion. However, expansions were less frequent than deletions, suggesting they may be distinct outcomes of a process, in particular, DNA polymerase slippage, rather than sister chromatid exchange (Gragg et al. 2002). PolydG::polydC tracts were also substantiated by independent sequencings but appeared to be less frequent than polydA::polydT tracts. Deletions are not biased toward the longest tracts, constituting only a small subset of the 215 tracts and the 55 tracts present in the genome for polydA and polydG tracts, respectively. Hence, a process of polymerase slippage is either restricted in frequency or acts in a site-dependent manner. We note that homopolymeric tracts themselves are not dispersed uniformly, with entire chromosomes missing either the polydA::polydT (chromosome 7) or polydG::polydC (chromosome 8) tracts, suggesting either different selective pressures or evolutionary sources of these sequences.

We also demonstrated the utility of Deep Variant for tracking down mutations likely responsible for phenotypes observed during growth. The adventitious mycelial phenotype that occurred between G5 and G10 was identified as a unique single mutation in the putative *flbA* gene (previously unidentified), by comparing SNP presence or absence at different growth stages. The studies coupled with CDD and AlphaFold3 analysis identified a putative FlbA structural homolog. This is a simple and rapid approach that can provide a more focused direction for targeted genetic analysis in organisms that are not amenable to genetic screens or selection, but for which real time data on the appearance of a phenotypic defect is observed.

In summary, these studies provide an overview of the genetic landscape of spontaneous events across the *A. flavus* genome, which is a foundational tool for future functional studies. We also present novel applications of long-read sequencing to analyze elusive biological processes.

## MATERIALS AND METHODS

### Strains, media, and growth conditions

Strain NRRL3357 was used for all of these studies. Gene identifiers following the AFLA and F9C07 nomenclatures are derived from A. flavus NRRL3357 reference assemblies JCVI-afl1-v.2.0 (GeneBank GCA_000006275.2and ASM901741v1 (GeneBank GCA_009171418.1), respectively (Skerker et al. 2021b; Fountain et al. 2020).

Mycelia of the strain NRRL3357 were vegetatively grown in serial weeklong cycles G5 (five weeks), G10 (ten cycles), and G15 (fifteen cycles) and DNA isolated as described previously (Chang et al. 1993; Adye and Mateles 1964; Woods et al. 2026), using the Qiagen Plant Mini kit (Qiagen, Beverly, MA) following the manufacturer’s protocol. Lyophilized mycelia were ground to a fine powder in liquid nitrogen and stored at −80 °C until needed. Briefly, 15 mg of each ground sample was processed in duplicate. After a spin column, the DNA was eluted with two 100 µl elutions using Qiagen AE buffer. The DNA suspension was further enriched by phenol/chloroform/isoamyl alcohol extraction and ethanol precipitation as described (Woods et al. 2026). DNA quantity and quality were determined using a NanoDrop OneC (Thermo Scientific). The quality of genomic DNA was also determined by electrophoresis on a 0.7% agarose gel.

### Quantification of asexual spore production

Quantification of asexual spore production in the *A. flavus* NRRL3357 wild type and GEN15 mutant was performed as described (Lohmar et al. 2025) with minor modifications. Briefly, the strains were single point inoculated onto 25 mL of GMM (Kafer 1977) and grown under light conditions for 7 days at 30°C. Conidiospores were harvested into sterile water, quantified utilizing an Olympus Automated Cell Counter Model R1 (Olympus Corporation, Shinjuku, Tokyo, Japan), and further adjusted to a concentration of 1.0 x 10^6^ spores/mL. Approximately 2 µL of spore suspension was single point inoculated onto 25 mL of Potato Dextrose Agar (BD Difco, Franklin Lakes, NJ, USA), YGT (Lohmar et al. 2019), Wickerham (Chang et al. 2012), and GMM media in triplicate. The cultures were incubated under light and dark conditions at 30°C for 7 days. After incubation, the cultures were photographed and a 7 mm core was taken approximately 1 cm away from the center of each culture. The cores were homogenized in 500 µL of sterile water and conidiospores were quantified as previously described. Statistical analysis was carried out for all quantitative data using the program R version x64 4.4.2 https://cran.r-project.org/manuals.html. Analysis of Variance (ANOVA) in conjunction with Games-Howell test. Since the variance of wild type and mutant cells were significantly different, the Tukeys test, in which the total variance of the dataset is distributed among all the samples was not an appropriate method. Data was considered statistically different with a p-value ≤ 0.05.

### Bioinformatic-Alpha Fold, CDD, homology comparison

Blast P and C-terminal Domain Database were searched by standard techniques. Protein structural similarity was measured by Alpha Fold 3 using the ChimeraX Matchmaker structural program. The root mean structural deviation (RMSD) was determined for each pair of protein structures.

### Sequencing Methodology

CD Genomics (Shirley, NY) carried out all sequencing. Both methods gave an average of 500-1500-fold coverage, although not equally distributed over all parts of the genome. ONT (v.10.4) was sequenced in a dual PromethION cell using 10.4 sequencing as described (Woods et al. 2026). PacBio HiFi sequencing library formation and sequencing methods were as previously described (Woods et al. 2026). Using the SMRTbell library was constructed using the SMRTbell Express Template Prep kit 2.0 (Pacific Biosciences).After polishing and tailing and ligation, the SMRTbell library was purified and subjected to a size selection step using the BluePippin system (Sage Science, Beverly, MA) to remove SMRTbells ≤15 kb. After size selection, the library was purified with 1X AMPure PB beads. The sequencing primer and Sequel II DNA Polymerase were annealed and bound, respectively, to the final SMRTbell library. The library was sequenced using 8M SMRT Cell on the Sequel II System with Sequel II Sequencing Kit (Hon et al. 2020).

### Assemblies

We bootstrapped G0 and G15 assemblies using both *A. flavus* strain NRRL3357 (GCA_009017415.1_ASM901741v1) (Skerker et al. 2021a) and mitochondrial (*A. flavus* SRRC1009) (Park et al. 2020) sequences that were concatenated to prepare a genome fasta file. This file served as input for G0 ONT or G15 PacBio reads to Rebaler (https://github.com/rrwick/Rebaler) (Wick et al. 2019). All assemblies for these studies have been reported previously (Woods et al. 2026).

### Structural variant determination

Structural variants were identified relative to the ONT G0 assembly using Sniffles 2 (version 2.2) (Smolka et al. 2024a). Alignments used minimap 2 (Li 2018) (version 2.26-r1175) with a gap open and gap extension set at 16 and 2, respectively. Structural variants were identified in algorithms as ≥ 20 bp deletions or expansions with variant read numbers (DV) of at least 2, as specified in the Results. This version was determined empirically to be only ∼70% efficient for low abundance sequences in simple sequence regions, leading to both false positives and false negatives. We therefore often coupled the algorithm with visual confirmation using IGV (version 2. (10.0)) (Thorvaldsdottir et al. 2013; Priyam et al. 2019). In the case of clustered and homopolymeric deletions, individual reads with copy numbers of six or less (or at least five random reads in cases of copy numbers of greater than six), were then aligned by BLAST analysis from these PacBio sequenced DNA to the G15 or G0 assemblies using SequenceServer (London, UK) (Priyam et al. 2019) as a step to eliminate areas of poor sequencing and to confirm the algorithmic call. We analyzed simple sequence organization for AT-rich sequences by removing the dust filter. For all studies, the variant frequency (VAF) calculated as (variant read number (DV)/reference read number (DR) x 100). In individual structural variants, VAF values ranged from 0.2 to 12.5 % (check) consistent with a high coverage. To ensure that clustered deletions defined were not the product of poorly sequenced regions, we made the conservative criteria that included only sequencing reads that were within 200 bp and did not contain a second deletion above 25% of the length of the original. Other sequences that were of overall poor quality were also eliminated by inspection.

IGV also revealed small errors (1-10 bp) in the G0 genomic assembly in the regions of polydG::polydC homopolymeric tracts that differed for each site of deletion, the presumed result of bootstrapping the assembly from a published dataset of 3357. This error was corrected by averaging the length of deletions that were present in reads that did not contain the large deletion and subtracting that estimate from the observed tract length.

### Pepper-Margin-Deep Variant Analyses of Single Nucleotide Polymorphisms Across Multiple Generations

Variant detection from PacBio Revio long reads was performed using the PEPPER-Margin-DeepVariant pipeline [(Shafin et al. 2021) v0.8; https://github.com/kishwarshafin/pepper] which integrates neural-network-based signal polishing (PEPPER), haplotype refinement (Margin) and variant calling (Deejavascript:;pVariant). To manage whole-genome variant calling in a controlled manner, the analysis was performed sequentially on a per-chromosome basis. For each chromosome, aligned reads were extracted from the full BAM file using Samtools v1.20 (https://www.htslib.org) to generate a chromosome-specific BAM file. A corresponding chromosome-restricted reference FASTA was also prepared, and all required inputs were organized into a structured working directory.

Variant calling was performed using the official kishwars/pepper_deepvariant:r0.8 Docker image. A custom wrapper script (runPEPPER-Margin-DeepVariant.sh), modeled directly from the command usage recommended in the official documentation, was used to launch the workflow with minor modifications to accommodate chromosome-specific inputs and output paths. The pipeline was run in HiFi mode (--hifi) with the chromosome-restricted BAM file and reference provided as inputs. This produced a set of compressed per-chromosome VCFs (*.vcf.gz) suitable for downstream filtering, annotation, and interpretation.

The variant analysis was compared to the ONT G0, G5, and G10 assemblies. The entire data set of single nucleotide polymorphisms was analyzed for those having high VAF values greater than 90% and present in G0 and G5 data sets but not in G10, since the conidial phenotype arises between G5 and G10. Hence variants would be expected in G0 and G5, but not in any subsequent dataset (G10 or G15). Only one single base pair mutation fulfilled these criteria. We also examined the Sniffles algorithm for translocations, inversions, and insertions in both genomic and mitochondrial sequences for any other potential association with the phenotype. but found none.

### Source of Homopolymer Data (Strains and Methods)

The sequencing data of a random set of ascomycetes fungi were derived from the strains under the conditions specified in Supplementary Table 2. The datasets are all available as the indicated assemblies at the NCBI genome site.

## DATA AVAILABILITY STATEMENT

## The authors affirm that all data necessary for confirming the conclusions of the article are present within the article, figures, tables, and supplementary data. . Strains and plasmids are available upon request. Sequencing datasets have been described (Woods et al. 2026)

## ACKNOWLEDGEMENTS

We thank Dr. Jeff W. Cary for his expert assistance and technical support in the early stages of these studies and Dr. Erik Flemington for his support. We also thank JWC and Bonnie Hoffman for critical review of the manuscript. AJL was funded by USDA NCA 58-6054-0-014. JML was funded by the USDA. The Next Generation Sequencing Analysis Core is supported by the National Institutes of Health National Cancer Institute, LSUHSC Tumor Virology COBRE, and the Louisiana Cancer Research Consortium.

Conceptualization: AW, JML and AJL; methodology: JML and AJL; investigation: AM, AW, JML and AJL; visualization: AW, JML and AJL; supervision: AJL; writing-original draft: AJL, writing-review and editing: JML, AW, and AJL.

The material is based upon work supported by the U.S. Department of Agriculture, Agriculture Research Service, under agreement NCA 58-6054-0-14. Any opinions, findings, conclusion, or recommendations expressed in this publication are those of the authors and do not necessarily reflect the view of the U.S. Department of Agriculture.

