## Supplemental Data for "The Genetic Landscape of Structural Variants in *Aspergillus flavus* using ONT and PacBio Sequencing"

Table of Contents:

| **Title** | Description | Page Number |
| --- | --- | --- |
| Figure S1 | Conidial Phenotypes | 1 |
| Figure S2 | BLASTP Analysis of flbA Homologs | 2 |
| Figure S3 | CDD Analysis of flbA Homologs | 3 |
| Figure S4 | Alpha Fold 3 Analysis of flbA Homologs | 3 |
| Figure S5A | Examples of 5 Deletions from A Site of PolydA::PolydT Homopolymeric Deletion | 5 |
| Figure S5B | Examples of 5 Deletions of from One Site of PolydG::PolydC Deletion | 6 |
| Table S1A | Centromeric Deletion Clusters | 8 |
| Table S1B | Subtelomeric/Other ATE Deletion Clusters | 9 |
| Table S1C | Dispersed Deletion Clusters | 10 |
| Table S2A | Coordinates, Genomic Context and Tract Length of Homopolymeric PolydA::PolydT Tracts. | 11 |
| Table S2B | Coordinates, Genomic Context and Tract Length of Homopolymeric PolydG::PolydC Tracts. | 17 |
| Table S3 | Tract Locations in Other Species | 19 |
| Table S4 | Species Accession Info | 24 |

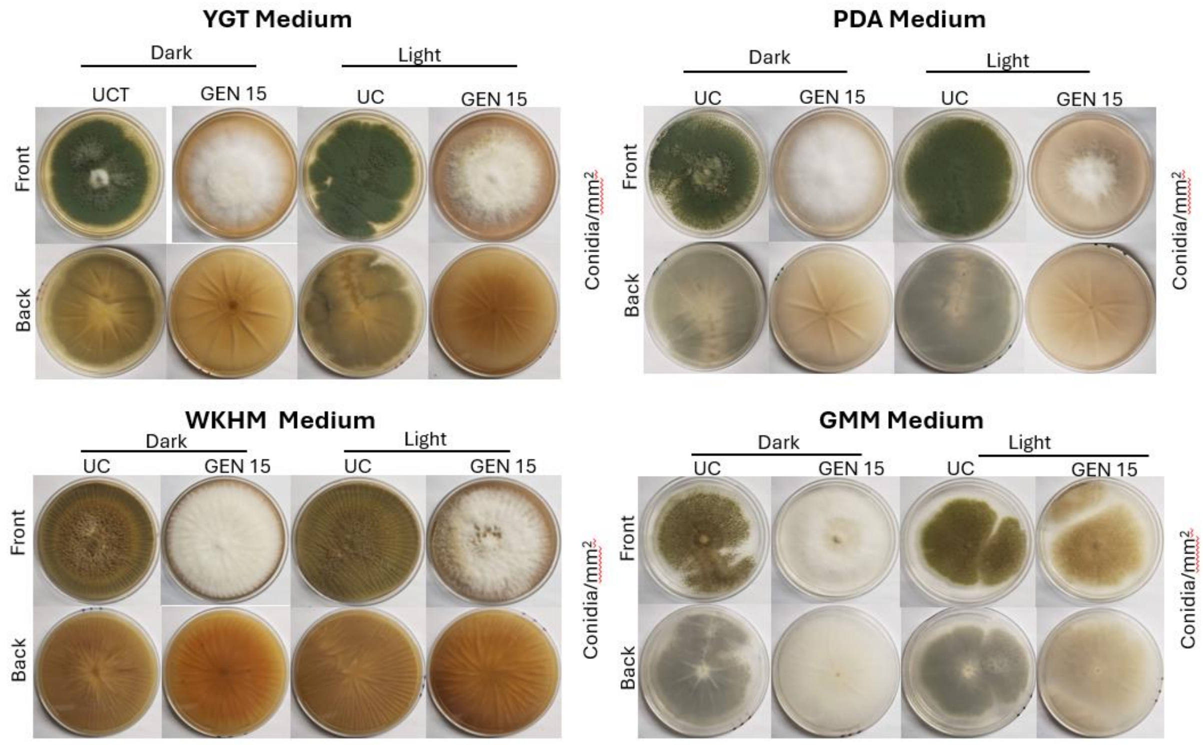

Figure S1: Full set of front and back of plates for the NRRL3357 wild type and G15 (labeled GEN15) mutant strain after growth on PDA, YGT, Wickerham (WKHM), and GMM media for 7 days under light and dark conditions at 30˚C highlighting the phenotypic differences in conidial formation.

```
Range 1: 1 to 719 GenPept  Graphics                     ▼ Next Match  ▲ Previou

Score           Expect Method                       Identities     Positives      Gaps
1117 bits(2888) 0.0    Compositional matrix adjust. 604/735(82%)   644/735(87%)   23/735(3%)

Query  1    MPTISSAHLDQDSPPSSHLLYS------QSPPSVDPIPVSTSPVETRPRTASAPSLQHNS  54
            MPTISS  LDQ SPPSSHL +S      QS  S  P+PVS+SP +    T  + S    S
Sbjct  1    MPTISSIPLDQGSPPSSHLRHSHHRLPTQSSLSNIPVPVSSSPDDFPSSTPPSASCTLTS  60

Query  55   APSAVVAPSQPATTTTTTTTTTTNNTPTGFTGSVIGSISRRNRRSFAALAREKTSNALAN  114
            A  AV AP+   +T T+TT+ T         GSVIGSISRRNRRSFAALAREKTSNA AN
Sbjct  61   ATPAVAAPTTRTSTFTSTTSFT---------GSVIGSISRRNRRSFAALAREKTSNAFAN  111

Query  115  LSSIGAP-NYPLRSSASSGSLSKHSRKASQVSVSEAPGATPLTPPLSDGSASSERSSPAP  173
            LS+IG   N  LRSSASSGSLSKHSRK SQVSVSEA GA+PLTPPLSD S SSE+ S  P
Sbjct  112  LSAIGGTTNATLRSSASSGSLSKHSRKLSQVSVSEATGASPLTPPLSDSSTSSEQLSNTP  171

Query  174  FDAATSRSSPAVVEQQTSSVERRRYTLQRAPSPLEQLPEVGPTVSPAKMHQTSSRLLRMT  233
             ++  +  +PAV EQ    VERRR TLQ  PS + Q  +    V+P+KMHQTSSRLLRMT
Sbjct  172  IESPANPITPAVAEQ----VERRRQTLQHVPSLVGQQAQA---VTPSKMHQTSSRLLRMT  224

Query  234  EDDRPFTKDFMDLFSTLMVSLKLDSHRVRFTKYDHSFTSEEAINNLGSLKFSQSNRMPDP  293
            EDDRPFTKDFMDLFSTLMVSLKLDSHRVRFT+YDHSFTSEEAINNLGSLKFSQSNRMPDP
Sbjct  225  EDDRPFTKDFMDLFSTLMVSLKLDSHRVRFTRYDHSFTSEEAINNLGSLKFSQSNRMPDP  284

Query  294  KDPSRIVTTTTTTTFSMAKEMARSVCQRFVDARFIEAVDGKSAQIFPLKGALYQLTPKGI  353
            KDPSRIVTTTTTTTFSMAKEMARSVCQRFVDARFIE+VDGK++Q FPLKGALYQLTPKGI
Sbjct  285  KDPSRIVTTTTTTTFSMAKEMARSVCQRFVDARFIESVDGKASQFFPLKGALYQLTPKGI  344

Query  354  NILQRFCQRNGITARHVIDVLESPRNTMQLVNLERDTDTDKLSHDRATIEVIFRRFAGQD  413
            NILQRFCQRNGITARHVIDVLESPRNTMQLVNLERD++TDKLSHDRATIEVIFRRFAGQD
Sbjct  345  NILQRFCQRNGITARHVIDVLESPRNTMQLVNLERDSETDKLSHDRATIEVIFRRFAGQD  404
```

Figure S2: BLASTP analysis of the putative flbA homolog (*AFLA_134030* ,query) compared to the Aspergillus nidulans flbA revealing 82% identity and 87% positives indicating a strong homology between the two species.

### A. Flavus CDD

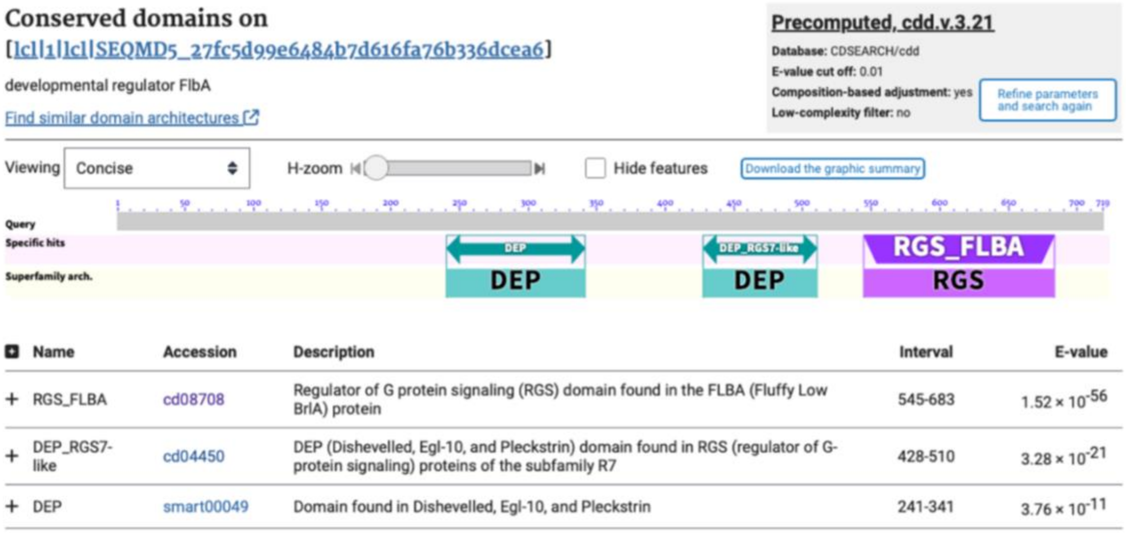

| Name | Accession | Description | Interval | E-value |
|---|---|---|---|---|
| + RGS_FLBA | cd08708 | Regulator of G protein signaling (RGS) domain found in the FLBA (Fluffy Low BrlA) protein | 545-683 | $1.52 \times 10^{-56}$ |
| + DEP_RGS7-like | cd04450 | DEP (Dishevelled, Egl-10, and Pleckstrin) domain found in RGS (regulator of G-protein signaling) proteins of the subfamily R7 | 428-510 | $3.28 \times 10^{-21}$ |
| + DEP | smart00049 | Domain found in Dishevelled, Egl-10, and Pleckstrin | 241-341 | $3.76 \times 10^{-11}$ |

Figure S3: CDD analysis of the putative flbA homlog (*AFLA_134030*; query) compared with the A. nidulans flbA species revealing strong homology with the RGS and DEP domains. The pflbA mutation lies within the conserved RGS domain.

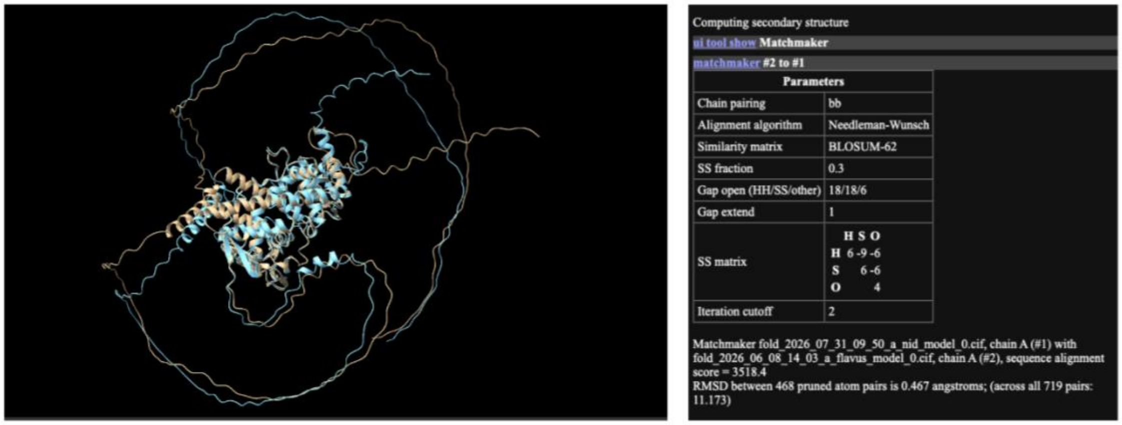

| Parameters | |
|---|---|
| Chain pairing | bb |
| Alignment algorithm | Needleman-Wunsch |
| Similarity matrix | BLOSUM-62 |
| SS fraction | 0.3 |
| Gap open (HH/SS/other) | 18/18/6 |
| Gap extend | 1 |
| SS matrix | H S O / H 6 -9 -6 / S 6 -6 / O 4 |
| Iteration cutoff | 2 |

Figure S4: AlphaFold 3 Analysis of Predicted Stabilities of the A. flavus pflbA (gold) with the A. nidulans flbA (blue) showing strong homology in the globular domain and an RMSD of 0.467 indicating alignment between the two proteins.

31bp

```
                ||||||||||||||||||||||||||||||||||||||||||||||||||||||||||||||||||||||||||||||||||||||||||||
Subject 5249811 CTCTTGTTCCCCTTTCTTATTCAAAAGAGTTTTATAATATCTCTTCTTAATAATGAGAAGGACGATTATAATTATACACCTTATGTGAAGCT 5249902

Query     11476 CATTTGATCGCCAACCTAGGGTGTGGAGGATTCTTCCTAACAGATCAATGATCACCGATGTGGTTGTATCATACTACGGAGTACTAGTAATA 11567
                ||||||||||||||||||||||||||||||||||||||||||||||||||||||||||||||||||||||||||||||||||||||||||||
Subject 5249903 CATTTGATCGCCAACCTAGGGTGTGGAGGATTCTTCCTAACAGATCAATGATCACCGATGTGGTTGTATCATACTACGGAGTACTAGTAATA 5249994

Query     11568 GTACGAATATTATTAATTGGTTGTTCCTTTTCTTTTTTC-------------------------------TTTCTTTTTTTTTTTTTTTTTT 11628
                |||||||||||||||||||||||||||||||||||||||                               ||| ||||||||||||||||||
Subject 5249995 GTACGAATATTATTAATTGGTTGTTCCTTTTCTTTTTTCTTTTCTTTTTTTTTTTTTTTTTTTTTTTTTTTTTTTTTTTTTTTTTTTTTTTTT 5250086

Query     11629 TTGTTTTTTTCCCTTTCTTTTTTTGCCTTAGGGGAAGGGCTGGTTGCAGTCTATCAGTCCACAGTGTGCAGGTACTGTACTTAGGGTACTGT 11720
                |||||||||||||||||||||||||||||||||||| |||||||||||||||||||||||||||||||||||||||||||||||||||||||
Subject 5250087 TTGTTTTTTTCCCTTTCTTTTTTTGCCTTAGGGGAA-GGCTGGTTGCAGTCTATCAGTCCACAGTGTGCAGGTACTGTACTTAGGGTACTGT 5250177

Query     11721 ACCCGTGACCTTACATACCTAGTGTGACCGACCCAAAGGTGGTACAGGCTCCATCAACTACGTTCTAGTCAGTCCCTCCTTGGTACTTATTT 11812
```

22bp

```
Subject 5249857 TTAATAATG-AGAAGGACGATTATAATTATACACCTTATGTGAAGCTCATTTGATCGCCAACCTAGGGTGTGGAGGATTCTTCCTAACAGAT 5249947

Query      7701 CAATGATCACCGATGTGGTTGTATCATACTACGGAGTACTAGTAATAGTACGAATATTATTAATTGGTTGTTCCTTTTCTTTTTTCTTTTCT 7792
                ||||||||||||||||||||||||||||||||||||||||||||||||||||||||||||||||||||||||||||||||||||||||||||
Subject 5249948 CAATGATCACCGATGTGGTTGTATCATACTACGGAGTACTAGTAATAGTACGAATATTATTAATTGGTTGTTCCTTTTCTTTTTTCTTTTCT 5250039

Query      7793 TTTTTTTTTTTTTTTTTTTTTTTTTTTTTTGTTTTT----------------------CCCTTCTTTTTTTGCCTTAGGGGAAGGCTGGTTG 7862
                |||||||||||||||||||||||||||||| |||||                      || |||||||||||||||||||||||||||||||
Subject 5250040 TTTTTTTTTTTTTTTTTTTTTTTTTTTTTTTTTTTTTTTTTTTTTTTTTGTTTTTTTCCCTTTCTTTTTTTGCCTTAGGGGAAGGCTGGTTG 5250131
```

19bps

```
Query      5795 TCACCGATGTGGTTGTATCATACTACGGAGTACTAGTAATAGTACGAATATTATTAATTGGTTGTTCCTTTTCTTTTTTCTTTTCTTTTTTT 5886
                ||||||||||||||||||||||||||||||||||||||||||||||||||||||||||||||||||||||||||||||||||||||||||||
Subject 5249954 TCACCGATGTGGTTGTATCATACTACGGAGTACTAGTAATAGTACGAATATTATTAATTGGTTGTTCCTTTTCTTTTTTCTTTTCTTTTTTT 5250045

Query      5887 TTTTTTTTTTTTTTTTTTTTTTTT-------------------GTTTTTTTCCCTTTCTTTTTTTGCCTTAGGGGAAGGCTGGTTGCAGTCT 5959
                ||||||||||||||||||||||||                   |||||||||||||||||||||||||||||||||||||||||||||||||
Subject 5250046 TTTTTTTTTTTTTTTTTTTTTTTTTTTTTTTTTTTTTTTTTTTGTTTTTTTCCCTTTCTTTTTTTGCCTTAGGGGAAGGCTGGTTGCAGTCT 5250137
```

37bp

```
Query     16646 ACCTGCACACTGTGGACTGATAGACTGCAACCAGCCTTCCCCTAAGGCAAAAAAAGAAAGGG------------------------------ 16707
                ||||||||||||||||||||||||||||||||||||||||||||||||||||||||||||||
Subject 5250158 ACCTGCACACTGTGGACTGATAGACTGCAACCAGCCTTCCCCTAAGGCAAAAAAAGAAAGGGAAAAAACAAAAAAAAAAAAAAAAAAAAAAA 5250067

Query     16708 -------AAAAAAAAAAAAAAAAAAAAAGAAAAGAAAAAAGAAAAGGAACAACCAATTAATAATATTCGTACTATTACTAGTACTCCGTAGT 16792
                       |||||||||||||||||||||||||||||||||||||||||||||||||||||||||||||||||||||||||||||||||||||
Subject 5250066 AAAAAAAAAAAAAAAAAAAAAAAAAAAAGAAAAGAAAAAAGAAAAGGAACAACCAATTAATAATATTCGTACTATTACTAGTACTCCGTAGT 5249975
```

39bp

```
Query      9568 GTACAGTACCTGCACACTGTGGACTGATAGACTGCAACCAGCCTTCCCCTAAGGCAAAAAAAGAAAGGGAAAAAACAAAAAAAAAAAA---- 9655
                ||||||||||||||||||||||||||||||||||||||||||||||||||||||||||||||||||||||||||||||||||||||||
Subject 5250165 GTACAGTACCTGCACACTGTGGACTGATAGACTGCAACCAGCCTTCCCCTAAGGCAAAAAAAGAAAGGGAAAAAACAAAAAAAAAAAAAAAA 5250074

Query      9656 -----------------------------------GAAAAGAAAAAAGAAAAGGAACAACCAATTAATAATATTCGTACTATTACTAGTACT 9712
                                                   |||||||||||||||||||||||||||||||||||||||||||||||||||||||||
Subject 5250073 AAAAAAAAAAAAAAAAAAAAAAAAAAAAAAAAAAAGAAAAGAAAAAAGAAAAGGAACAACCAATTAATAATATTCGTACTATTACTAGTACT 5249982
```

Supplementary Figure 5A: Examples of 5 deletions (highlighted in yellow)of indicated size in base pairs a site  of PolydA::polydT homopolymeric Deletion at Chromosome 1: coordinate **520072** (query) relative to the NRRL 3357 assembly (subject).

18bp

```
Query    20273 GAAACACTACTTGCAATCATGGCTACCGTGGTACCTAAATCTCCTTCCTCCTCCTTTCCT-------------- 20332
               ||||||||||||||||||||||||||||||||||||||||||||||||||||||||||||
Subject 185390 GAAACACTACTTGCAATCATGGCTACCGTGGTACCTAAATCTCCTTCCTCCTCCTTTCCTCCCCCCCCCCCCCC 185317
Query    20333 ----CCCCCCCCCCCCGGGCTACGGTTTAAACTAACTGGGTATACTAATGAATCAGCCCGAAGGCTGGTCCAAC 20402
                   ||||||||||||||||||||||||||||||||||||||||||||||||||||||||||||||||||||||
Subject 185316 CCCCCCCCCCCCCCCCGGGCTACGGTTTAAACTAACTGGGTATACTAATGAATCAGCCCGAAGGCTGGTCCAAC 185243
```

22bp

```
Query     9835 GGACTGTCGTCCGTCGACCAATACTCATCCAGATCACTTGGGTTGTTGGACCAGCCTTCGGGCTGATTCATTAG 9908
               ||||||||||||||||||||||||||||||||||||||||||||||||||||||||||||||||||||||||||
Subject 185199 GGACTGTCGTCCGTCGACCAATACTCATCCAGATCACTTGGGTTGTTGGACCAGCCTTCGGGCTGATTCATTAG 185272
Query     9909 TATACCCAGTTAGTTTAAACCGTAGCCCGGGGGGAG-----------------------GGAAAGGAGGAGGAA 9959
               |||||||||||||||||||||||||||||||||| |                       |||||||||||||||
Subject 185273 TATACCCAGTTAGTTTAAACCGTAGCCCGGGGGGGGGGGGGGGGGGGGGGGGGGGGGGAGGAAAGGAGGAGGAA 185346
Query     9960 GGAGATTTAGGTACCACGGTAGCCATGATTGCAAGTAGTGTTTCAAAGTTGTTGTTGTTGGAGATTGGCGG 10033
               ||||||||||||||||||||||||||||||||||||||||||||||||||||||||||||||||||||||||||
Subject 185347 GGAGATTTAGGTACCACGGTAGCCATGATTGCAAGTAGTGTTTCAAAGTTGTTGTTGTTGGAGATTGGCGG 185420
```

18bp

```
Query     6288 ACCCCCCGGATATCCTTTTCAGTCTCCGCCAATCTCCAACAACAACAACAACTTTGAAACACTACTTGCAATCA 6361
               ||||||||||||||||||||||||||||||||||||||||||||||||||||||||||||||||||||||||||
Subject 185445 ACCCCCCGGATATCCTTTTCAGTCTCCGCCAATCTCCAACAACAACAACAACTTTGAAACACTACTTGCAATCA 185372
Query     6362 TGGCTACCGTGGTACCTAAATCTCCTTCCTCCTCCTTTCCTCCCCCCCCCC--------------------GGG 6415
               |||||||||||||||||||||||||||||||||||||||||||||||||||                    |||
Subject 185371 TGGCTACCGTGGTACCTAAATCTCCTTCCTCCTCCTTTCCTCCCCCCCCCCCCCCCCCCCCCCCCCCCCCGGG 185298
Query     6416 CTACGGTTTAAACTAACTGGGTATACTAATGAATCAGCCCGAAGGCTGGTCCAACAACCCAAGTGATCTGGATG 6489
               ||||||||||||||||||||||||||||||||||||||||||||||||||||||||||||||||||||||||||
Subject 185297 CTACGGTTTAAACTAACTGGGTATACTAATGAATCAGCCCGAAGGCTGGTCCAACAACCCAAGTGATCTGGATG 185224
```

16bp

```
Query    14113 CCCCGGATATCCTTTTCAGTCTCCGCCAATCTCCAACAACAACAACAACTTTGAAACACTACTTGCATTCATGG 14186
               ||||||||||||||||||||||||||||||||||||||||||||||||||||||||||||||||||| ||||||
Subject 185442 CCCCGGATATCCTTTTCAGTCTCCGCCAATCTCCAACAACAACAACAACTTTGAAACACTACTTGCAATCATGG 185369
Query    14187 CTACCGTGGTACCTAAATCTCCTTCCTCCCTCCTT-----------------CCCCCCCCCCCCCCCCCGGGCT 14243
               ||||||||||||||||||||||||||||| |||||                 ||||||||||||||||||||||
Subject 185368 CTACCGTGGTACCTAAATCTCCTTCCTCC-TCCTTTCCTCCCCCCCCCCCCCCCCCCCCCCCCCCCCCGGGCT 185296
Query    14244 ACGGTTTAAACTAACTGGGTATACTAATGAATCAGCCCGAAGGCTGGTCCAACAACCCAAGTGATCTGGATGAG 14317
               ||||||||||||||||||||||||||||||||||||||||||||||||||||||||||||||||||||||||||
Subject 185295 ACGGTTTAAACTAACTGGGTATACTAATGAATCAGCCCGAAGGCTGGTCCAACAACCCAAGTGATCTGGATGAG 185222
```

28bp

```
Query     6213 AATCTCCAACAACAACAACAACTTTGAAACACTACTTGCAATCATGGCTACCGTGGTACCTAAATCTCCTTCCT 6286
               ||||||||||||||||||||||||||||||||||||||||||||||||||||||||||||||||||||||||||
Subject 185415 AATCTCCAACAACAACAACAACTTTGAAACACTACTTGCAATCATGGCTACCGTGGTACCTAAATCTCCTTCCT 185342
Query     6287 CCTCCTTTCC---------------------------CCCCGGGCTACGGTTTAAACTAACTGGGTATACTAAT 6333
               ||||||||||                           |||||||||||||||||||||||||||||||||||||
Subject 185341 CCTCCTTTCCTCCCCCCCCCCCCCCCCCCCCCCCCCCCCCCGGGCTACGGTTTAAACTAACTGGGTATACTAAT 185268
Query     6334 GAATCAGCCCGAAGGCTGGTCCAACAACCCAAGTGATCTGGATGAGTATTGGTCGACGGACGACAGTCCTGGAA 6407
               ||||||||||||||||||||||||||||||||||||||||||||||||||||||||||||||||||||||||||
Subject 185267 GAATCAGCCCGAAGGCTGGTCCAACAACCCAAGTGATCTGGATGAGTATTGGTCGACGGACGACAGTCCTGGAA 185194
```

Supplementary Figure 5B: Examples of 5 Deletions (highlighted in yellow) of indicated size in base pairs from a site of PolydG::PolydC homopolymeric deletion at chromosome 1: **185303**(query) relative to the NRRL 3357 assembly (subject).

| Chromosome | Coordinate (reads) | Deletion (bp) | Expected% AT | Actual % AT |
|---|---|---|---|---|
| **1** | 4540921 (2) | 20-24 | 90 | 93.5 |
| | 4598096 (3) | 18-30 | 90 | 97 |
| | 4598907 (2) | 22-24 | 90 | 96.5 |
| | 4600184 (4) | 20-23 | 90 | 94.5 |
| | 4610482 (2) | 19-30 | 90 | 96.5 |
| **2** | 2053014 (2) | 19-22 | 90 | 97.5 |
| | 2073497(2) | 18-27 | 90 | 95 |
| | 2097600 (2) | 22-23 | 90 | 94.5 |
| | 2112914 (2) | 24-48 | 90 | 95.5 |
| **3** | 2749003 (2) | 25-27 | 90 | 94.5 |
| | 2763105 (2) | 16-29 | 90 | 97 |
| | 2776651 (2) | 23-26 | 90 | 96.5 |
| | 2801654 (4) | 21-33 | 90 | 92.5 |
| | 2853652 (2) | 20-41 | 90 | 93.5 |
| **4** | 2733546 (2) | 18-25 | 90 | 94 |
| | 2804749 (2) | 19-29 | 90 | 97 |
| | 2824135 (2) | 20-23 | 90 | 92.5 |
| **5** | 2403550 (2) | 19-27 | 90 | 98 |
| | 2420879 (2) | 21-24 | 90 | 90.5 |
| | 2454561 (2) | 21-26 | 90 | 94 |
| **6** | 2115856 (2) | 24-30 | 90 | 94 |
| **7** | 2598793 (2) | 21-28 | 90 | 94.5 |
| | 2663333 (2) | 21-22 | 90 | 95 |
| | 2574310 (2) | 18-23 | 90 | 95.5 |
| | 2640233 (2) | 19-23 | 90 | 92 |
| | 2665191 (3) | 18-30 | 90 | 96 |
| **8** | 1852958 (2) | 18-21 | 90 | 96.5 |
| | 1854765 (2) | 19-22 | 90 | 90 |
| | 1858763 (2) | 19-22 | 90 | 95 |
| | 1877154 (2) | 21-24 | 90 | 93 |
| **Average** | 30 | 23.3 | 90 | 94.7 |

Table S1A: Primary Data for Table 2A compilation showing each centromeric deletion clustersas described in Table 2A.

| Chromosome | Coordinate | Deletion (bp) | Expected % AT | Actual % AT |
|---|---|---|---|---|
| **1** | 32579 (2) | 20-21 | 90 | 94.12 |
| | 34550 (2) | 22-30 | 90 | 95.77 |
| | 6195368 (2) | 23-42 | 86 | 95.78 |
| | 6509236 (2) | 18-41 | 85 | 95.04 |
| **2** | 336810 (3) | 20-40 | 90 | 97.29 |
| | 412344 (2) | 22-23 | 89 | 89.29 |
| | 3673862 (2) | 16-21 | 95 | 99 |
| **4** | 4809960 (2) | 24-28 | 89 | 79.32 |
| **5** | 3970 (2) | 15-33 | 92 | 96.3 |
| | 4545506 (2) | 19-20 | 90 | 91.86 |
| | 4548523 (2) | 16-23 | 90 | 91.71 |
| **7** | 688262 (2) | 18-24 | 93 | 92.69 |
| **8** | 2579748 (3 | 18-30 | 92 | 93.94 |
| **Average** | 13 | 24 | 90.1 | 93.3 |

Table S1B: Primary Data for Table 2B compilation showing each subtelomeric/other ATE deletion cluster as described in Table 2B.

| Chromosome | Chromosome | Deletion (bp) |
|---|---|---|
| **1** | 1101537 (2) | 20-24 |
| | 1712138 (2) | 19-40 |
| | 2165131 (2) | 19-28 |
| | 2594069 (2) | 19-20 |
| | 4462563 (2) | 20-34 |
| | 5505718 (2) | 24-29 |
| **2** | 2734117 (2) | 18-31 |
| **3** | 477836 (2) | 26-35 |
| | 1254670 (2) | 19-31 |
| | 1339705 (2) | 29-32 |
| | 1990832 (2) | 23-58 |
| **4** | 120839 (2) | 37-39 |
| | 1092626 (2) | 25-26 |
| | 2002903 (2) | 23-35 |
| | 2862603 (2) | 33-38 |
| | 3411126 (2) | 24-26 |
| **5** | 1674391 (2) | 19-45 |
| **6** | 398296 (2) | 65-88 |

| | | |
|---|---|---|
| | 991799 (2) | 17-19 |
| | 2551875 (2) | 22-33 |
| | 3061023 (2) | 42-51 |
| | 3203132 (2 | 20-22 |
| | 3858471 (2) | 23-60 |
| **7** | 756846 (2) | 49-50 |
| **8** | 887558 (2) | 24-39 |
| | 1008084 (2) | 26-67 |
| | 1292211 (2) | 20-22 |
| | 2575274 (2) | 23-54 |
| | 3129107 (2) | 21-22 |
| **Average** | 28 | 31.8 |

Table S1C: Primary Data for Table 2C compilation showing each dispersed deletion cluster as described in Table 2C.

| Chromosome | Coordinate | Coding | Feature | Tract Size |
|---|---|---|---|---|
| 1 | 4815870 | nc | | 60 |
| 1 | 5250063 | nc | | 60 |
| 1 | 1959115 | nc | | 59 |
| 1 | 4079501 | nc | | 54 |
| 1 | 3008144 | intron | | 47 |
| 1 | 5296604 | nc | | 43 |
| 1 | 5464487 | nc | | 43 |
| 1 | 1565507 | nc | | 41 |
| 1 | 3119800 | nc | | 41 |
| 1 | 6419678 | nc | | 40 |
| 1 | 2022643 | nc | | 39 |
| 1 | 4833627 | nc | | 39 |
| 1 | 5426223 | intron | | 39 |
| 1 | 5331515 | nc | | 37 |
| 1 | 5585786 | nc | | 36 |
| 1 | 4681866 | nc | | 34 |
| 1 | 395512 | nc | | 30 |
| 1 | 1366737 | nc | 3' | 30 |
| 1 | 3039325 | nc | | 30 |
| 1 | 2473094 | nc | | 29 |
| 1 | 3703573 | intron | | 29 |
| 1 | 379830 | nc | | 28 |
| 1 | 2001104 | nc | | 28 |
| 1 | 2924850 | nc | | 28 |
| 1 | 5179766 | nc | | 28 |
| 1 | 3173414 | nc | | 26 |
| 1 | 4761060 | nc | | 26 |
| 1 | 300794 | intron | | 25 |
| 1 | 2110921 | nc | | 24 |
| 1 | 2845435 | nc | | 24 |
| 1 | 4633211 | nc | | 24 |
| 1 | 3005311 | nc | | 23 |
| 1 | 3733268 | nc | 3' | 22 |
| 2 | 1032759 | intron | | 59 |
| 2 | 3210941 | nc | | 53 |
| 2 | 2426547 | nc | | 46 |
| 2 | 5198493 | nc | | 43 |

| | | | | | |
|---|---|---|---|---|---|
| **2** | 4095361 | | intron | | 40 |
| **2** | 4135281 | | nc | | 38 |
| **2** | 4138074 | | nc | 3' | 38 |
| **2** | 5129993 | | intron | | 38 |
| **2** | 2321724 | | nc | | 37 |
| **2** | 2935839 | | nc | 3' | 36 |
| **2** | 5528477 | | nc | | 35 |
| **2** | 2261055 | | nc | | 30 |
| **2** | 3448702 | | nc | | 30 |
| **2** | 2936261 | | nc | | 29 |
| **2** | 4442917 | | nc | | 29 |
| **2** | 6030526 | | nc | | 29 |
| **2** | 105515 | | nc | | 28 |
| **2** | 1979432 | | nc | | 28 |
| **2** | 2269304 | | nc | | 28 |
| **2** | 173551 | | intron | | 27 |
| **2** | 5343108 | | nc | 5' | 26 |
| **2** | 1577556 | | nc | | 25 |
| **2** | 2439225 | | nc | 5' | 25 |
| **2** | 1904453 | | nc | | 24 |
| **2** | 2588445 | | nc | | 24 |
| **2** | 1419349 | | nc | | 23 |
| **2** | 1624155 | | nc | | 23 |
| **2** | 5028324 | | nc | | 22 |
| **3** | 592961 | | nc | | 73 |
| **3** | 4843136 | | nc | | 49 |
| **3** | 2551031 | | intron | | 46 |
| **3** | 4635121 | | nc | | 42 |
| **3** | 1947483 | | nc | 3' | 41 |
| **3** | 5129154 | | nc | | 40 |
| **3** | 374016 | | nc | | 35 |
| **3** | 1367171 | | nc | | 34 |
| **3** | 43232 | | nc | | 30 |
| **3** | 1129434 | | nc | | 30 |
| **3** | 2607141 | | nc | | 29 |
| **3** | 3517600 | | nc | | 29 |
| **3** | 327073 | | nc | | 28 |
| **3** | 862277 | | nc | | 28 |
| **3** | 1159573 | | nc | | 28 |

| | | | | | |
|---|---|---|---|---|---|
| **3** | 4032475 | | nc | 3' | 28 |
| **3** | 71726 | | nc | | 27 |
| **3** | 239268 | | nc | 5' | 27 |
| **3** | 2162759 | | nc | | 27 |
| **3** | 2604651 | | nc | | 27 |
| **3** | 4117366 | | nc | | 27 |
| **3** | 584984 | | nc | | 26 |
| **3** | 1349404 | | nc | | 26 |
| **3** | 2500202 | | nc | | 26 |
| **3** | 3636608 | | nc | | 26 |
| **3** | 4085321 | | nc | | 26 |
| **3** | 1192084 | | nc | | 25 |
| **3** | 4241676 | | intron | | 25 |
| **3** | 4378793 | | nc | | 25 |
| **3** | 2485114 | | nc | | 24 |
| **3** | 2977988 | | nc | | 24 |
| **3** | 3642236 | | nc | | 23 |
| **3** | 3823388 | | nc | | 23 |
| **3** | 3215940 | | nc | | 22 |
| **4** | 3535411 | | nc | | 44 |
| **4** | 2898879 | | intron | | 41 |
| **4** | 3187560 | | nc | | 39 |
| **4** | 4431244 | | nc | | 39 |
| **4** | 490173 | | nc | | 38 |
| **4** | 3444814 | | intron | | 37 |
| **4** | 347764 | | nc | 3' | 36 |
| **4** | 2209854 | | nc | | 35 |
| **4** | 467815 | | nc | | 32 |
| **4** | 3150103 | | nc | | 32 |
| **4** | 3304725 | | nc | | 32 |
| **4** | 485715 | | nc | | 31 |
| **4** | 1202583 | | nc | 5' | 31 |
| **4** | 2673614 | | nc | | 31 |
| **4** | 4472279 | | nc | | 31 |
| **4** | 187443 | | nc | | 30 |
| **4** | 3898552 | | nc | | 30 |
| **4** | 3731614 | | nc | | 29 |
| **4** | 4535954 | | nc | | 29 |
| **4** | 1714702 | | nc | | 28 |

| | | | | | |
|---|---|---|---|---|---|
| **4** | 1329561 | | nc | 3' | 26 |
| **4** | 1474834 | | nc | 3' | 26 |
| **4** | 1572093 | | nc | 5' | 26 |
| **4** | 1482103 | | nc | | 25 |
| **4** | 4059885 | | nc | | 25 |
| **4** | 2643197 | | nc | | 24 |
| **4** | 2984172 | | nc | 3' | 24 |
| **4** | 3037270 | | nc | 3' | 23 |
| **4** | 3038269 | | nc | | 23 |
| **4** | 3267820 | | nc | | 23 |
| **4** | 2671827 | | nc | | 22 |
| **5** | 3715764 | | nc | | 45 |
| **5** | 3960053 | | nc | | 44 |
| **5** | 4155120 | | nc | | 44 |
| **5** | 4373383 | | nc | | 44 |
| **5** | 1242692 | | nc | 3' | 43 |
| **5** | 3099896 | | nc | | 39 |
| **5** | 3522725 | | nc | | 39 |
| **5** | 289049 | | nc | | 36 |
| **5** | 3424678 | | nc | | 35 |
| **5** | 2263474 | | nc | | 34 |
| **5** | 3183368 | | nc | | 34 |
| **5** | 2211453 | | nc | | 33 |
| **5** | 2953868 | | nc | | 33 |
| **5** | 3975630 | | nc | | 33 |
| **5** | 754308 | | nc | | 31 |
| **5** | 1946755 | | nc | | 31 |
| **5** | 1339432 | | nc | | 29 |
| **5** | 3116314 | | nc | | 29 |
| **5** | 3536239 | | nc | | 29 |
| **5** | 253065 | | intron | | 28 |
| **5** | 1793070 | | nc | | 28 |
| **5** | 2260477 | | nc | | 28 |
| **5** | 2159703 | | nc | | 27 |
| **5** | 3023925 | | nc | | 27 |
| **5** | 3793791 | | intron | | 27 |
| **5** | 3867269 | | nc | | 27 |
| **5** | 1008706 | | nc | | 26 |
| **5** | 457527 | | nc | | 25 |

| | | | | | |
|---|---|---|---|---|---|
| **5** | 581257 | | nc | | 25 |
| **5** | 2584061 | | nc | | 24 |
| **5** | 3624860 | | nc | | 24 |
| **6** | 3274393 | | nc | | 42 |
| **6** | 387347 | | nc | | 41 |
| **6** | 1746504 | | nc | | 41 |
| **6** | 2475405 | | nc | | 41 |
| **6** | 2032489 | | nc | | 39 |
| **6** | 428776 | | nc | | 38 |
| **6** | 891788 | | nc | | 38 |
| **6** | 3310503 | | nc | | 37 |
| **6** | 935894 | | nc | | 34 |
| **6** | 794296 | | nc | 3' | 33 |
| **6** | 1443817 | | nc | | 33 |
| **6** | 1544109 | | nc | | 33 |
| **6** | 1858987 | | nc | | 33 |
| **6** | 1967530 | | nc | | 32 |
| **6** | 3341134 | | nc | | 31 |
| **6** | 3297274 | | nc | | 29 |
| **6** | 897742 | | nc | | 28 |
| **6** | 1604261 | | nc | | 28 |
| **6** | 1658925 | | nc | | 28 |
| **6** | 3252578 | | nc | | 28 |
| **6** | 2451492 | | nc | | 27 |
| **6** | 468414 | | nc | | 26 |
| **6** | 530616 | | nc | | 26 |
| **6** | 2562491 | | nc | | 26 |
| **6** | 3816310 | | nc | | 26 |
| **6** | 2482344 | | nc | | 25 |
| **6** | 3743598 | | nc | | 25 |
| **6** | 602833 | | nc | | 24 |
| **6** | 1418825 | | nc | | 23 |
| **6** | 1085738 | | nc | | 22 |
| **8** | 660991 | | nc | | 52 |
| **8** | 1342333 | | intron | | 52 |
| **8** | 3095989 | | intron | | 49 |
| **8** | 1550755 | | nc | | 48 |
| **8** | 3010368 | | nc | 3' | 48 |
| **8** | 1461129 | | nc | | 43 |

| | | | | | |
|---|---|---|---|---|---|
| **8** | 1100888 | | nc | | 42 |
| **8** | 1216195 | | nc | | 42 |
| **8** | 2855106 | | nc | | 42 |
| **8** | 2726028 | | nc | | 41 |
| **8** | 1766278 | | intron | | 40 |
| **8** | 2622811 | | intron | | 32 |
| **8** | 1361463 | | nc | | 31 |
| **8** | 669681 | | intron | | 28 |
| **8** | 2100253 | | nc | | 28 |
| **8** | 908803 | | nc | | 27 |
| **8** | 2057054 | | nc | | 25 |
| **8** | 2717621 | | nc | | 25 |
| **8** | 59799 | | nc | 3' | 24 |
| **8** | 692154 | | intron | | 24 |
| **8** | 900052 | | nc | | 24 |
| **8** | 1977947 | | nc | | 24 |
| **8** | 2387012 | | nc | | 24 |
| **8** | 1290610 | | nc | | 23 |
| **8** | 3114377 | | nc | | 23 |
| **8** | 3158703 | | nc | | 23 |

Table S2A: Coordinates, genomic context and tract Length of homopolymeric polydA::polydT Tracts. Definitions: nc, non-coding; red text, sites identified as deletions or expansions; 5' and 3' refer to non-coding sequences 5' or 3' to the predicted direction of transcription.

| Chromosome | Coordinate | | Coding | Feature | Tract Size |
|---|---|---|---|---|---|
| **1** | 4156442 | | nc | 3' | 30 |
| **1** | 185303 | | nc | 5' | 29 |
| **1** | 4273135 | | nc | 5' | 29 |
| **1** | 4430959 | | nc | | 28 |
| **1** | 4034216 | | nc | | 27 |
| **1** | 4692189 | | nc | 5' | 26 |
| **1** | 1607707 | | nc | | 25 |
| **1** | 5321526 | | nc | | 25 |
| **1** | 5847328 | | nc | | 24 |
| **1** | 852655 | | nc | | 23 |
| **1** | 2215576 | | nc | | 22 |
| **1** | 5209736 | | nc | | 22 |
| **1** | 5326699 | | nc | | 22 |
| **1** | 5339845 | | nc | | 22 |
| **2** | 2263616 | | nc | | 34 |
| **2** | 3199731 | | nc | | 31 |
| **2** | 366254 | | nc | | 29 |
| **2** | 5766028 | | nc | | 29 |
| **2** | 2021840 | | intron | | 26 |
| **2** | 1954019 | | nc | | 25 |
| **2** | 2189182 | | intron | | 25 |
| **2** | 1899488 | | nc | | 24 |
| **2** | 3445745 | | nc | | 22 |
| **2** | 2316319 | | nc | | 20 |
| **3** | 3432056 | | nc | 3' | 48 |
| **3** | 4973080 | | nc | | 31 |
| **3** | 1899254 | | nc | | 24 |
| **3** | 2152438 | | nc | | 24 |
| **3** | 3109586 | | nc | 5' | 24 |
| **3** | 1846112 | | nc | | 23 |
| **3** | 2872740 | | nc | | 23 |
| **3** | 3842500 | | nc | | 20 |
| **4** | 1794375 | | intron | | 27 |
| **4** | 3237354 | | nc | | 24 |
| **4** | 1522414 | | nc | | 22 |
| **4** | 3794685 | | nc | | 22 |
| **5** | 3609504 | | nc | | 30 |

| | | | | | |
|---|---|---|---|---|---|
| **5** | 1997198 | | nc | | 27 |
| **5** | 1000605 | | intron | | 26 |
| **5** | 3389638 | | nc | | 26 |
| **5** | 3342672 | | nc | | 25 |
| **5** | 3229045 | | nc | | 24 |
| **5** | 259351 | | intron | | 21 |
| **5** | 442235 | | nc | 5' | 20 |
| **6** | 991799 | | nc | | 28 |
| **6** | 603965 | | nc | | 27 |
| **6** | 533201 | | nc | | 26 |
| **6** | 2003601 | | nc | | 25 |
| **6** | 212092 | | nc | | 22 |
| **7** | 1449018 | | nc | | 25 |
| **7** | 2694782 | | nc | | 25 |
| **7** | 1116056 | | nc | | 24 |
| **7** | 1730696 | | nc | | 23 |

Table S2B: Coordinates, genomic context and tract length of Homopolymeric polydG::polydC tracts. Definitions: nc, non-coding; red text, identified as deletions or expansions; 5' and 3' refer to non-coding sequences 5' or 3' to the predicted direction of transcription.

| ***Pyricularia oryzae Br48*** | | | **Candidozyma auris** | | | **Trichoderma reesei QM6a** | | | **Monascus purpureus KUPM5** | | | **Magnusiomyces sp. LA-1** | | |
|---|---|---|---|---|---|---|---|---|---|---|---|---|---|---|
| *Chr:* | Coor | bp | Chr: | Coor | bp | Chr: | Coor | bp | Chr | Coor | bp | Chr | Coor | bp |
| *1* | 5259517 | 60 | 1 | 61531 | 44 | 1 | 2054796 | 51 | 1 | 123920 | 61 | 1 | | |
| | 3139425 | 58 | | 129766 | 44 | | 4811028 | 53 | | 1410289 | 45 | | 3151 | 49 |
| | 3007388 | 56 | | 609331 | 56 | | 4869122 | 50 | 2 | 1652423 | 44 | | 123488 | 59 |
| | 51228 | 55 | | 630381 | 45 | | 5839031 | 45 | 3 | 0 | NA | | 133836 | 78 |
| | 5334598 | 55 | | 663651 | 57 | | 6887764 | 58 | 4 | 0 | NA | | 145447 | 48 |
| | 3809216 | 52 | | 882361 | 48 | 2 | 4058718 | 63 | 5 | 0 | NA | | 300350 | 49 |
| | 2520409 | 50 | | 994437 | 50 | | 5173420 | 52 | 6 | 0 | NA | | 397474 | 44 |
| | 2971352 | 49 | | 1044653 | 50 | | 321875 | 51 | 7 | 2100988 | 53 | | 453026 | 49 |
| | 3790665 | 49 | | 1063387 | 48 | 3 | 1520092 | 54 | 8 | 1245773 | 48 | | 469792 | 91 |
| | 4606862 | 49 | | 1070970 | 47 | 4 | 2948654 | 45 | | | | | 475385 | 48 |
| | 4578146 | 48 | | 1383412 | 64 | | 3561314 | 44 | | Total | 5 | | 487310 | 85 |
| | 5397396 | 48 | | 1544234 | 68 | 5 | 775719 | 49 | | | | | 526462 | 57 |
| | 5342989 | 47 | | 1578701 | 47 | | 1858651 | 44 | | | | | 531452 | 46 |
| | 1530658 | 46 | | 1592792 | 49 | | 2094989 | 55 | | | | | 566480 | 51 |
| | 3646918 | 46 | | 1849713 | 60 | | 3336980 | 44 | | | | | 573799 | 74 |
| | 2626423 | 45 | | 1976815 | 45 | | 3407336 | 54 | | | | | 618322 | 66 |
| | 4792547 | 45 | | 2014834 | 49 | | 3534102 | 55 | | | | | 657895 | 47 |
| | 4949016 | 45 | | 2133692 | 55 | | 4158612 | 51 | | | | | 720147 | 58 |
| | 5594115 | 45 | | 2595508 | 51 | 6 | 667176 | 55 | | | | | 727499 | 55 |
| | 2279279 | 44 | | 2640191 | 44 | 7 | 1108877 | 53 | | | | | 763159 | 67 |
| | 2397704 | 44 | | 2675256 | 54 | 8 | 0 | NA | | | | | 799213 | 57 |
| | 3821149 | 44 | | 2680600 | 45 | | | | | | | | 820827 | 65 |
| | 4061336 | 44 | | 2958135 | 45 | | Total | 20 | | | | | 1003171 | 48 |
| | 4687806 | 44 | | 3244679 | 76 | | | | | | | | 1033143 | 57 |
| *2* | 125550 | 48 | | 3428102 | 58 | | | | | | | | 1096060 | 60 |
| | 212061 | 47 | | 3535502 | 47 | | | | | | | | 1220778 | 82 |
| | 498639 | 53 | | 3559041 | 52 | | | | | | | | 1253970 | 58 |
| | 778184 | 47 | | 3870469 | 49 | | | | | | | | 1329104 | 56 |
| | 1166131 | 51 | | 3956809 | 53 | | | | | | | | 1338472 | 46 |
| | 1166715 | 44 | | 3979482 | 45 | | | | | | | | 1423395 | 53 |
| | 1856949 | 46 | 2 | 41466 | 45 | | | | | | | | 1443049 | 47 |
| | 1856956 | 46 | | 66569 | 44 | | | | | | | | 1443049 | 47 |
| | 2329524 | 53 | | 88382 | 49 | | | | | | | | 1510621 | 58 |
| | 2355714 | 45 | | 120413 | 51 | | | | | | | | 1518469 | 60 |
| | 2583409 | 46 | | 167527 | 75 | | | | | | | | 1548939 | 61 |
| | 2988330 | 52 | | 390642 | 53 | | | | | | | | 1566345 | 51 |
| | 5059110 | 46 | | 417645 | 50 | | | | | | | | 1703343 | 51 |
| | 5147117 | 46 | | 477507 | 72 | | | | | | | | 1757604 | 45 |
| | 5261897 | 46 | | 481745 | 46 | | | | | | | | 1874043 | 85 |
| | 5495003 | 51 | | 481997 | 45 | | | | | | | | 1962750 | 70 |
| | 5495013 | 51 | | 550380 | 51 | | | | | | | | 2055505 | 64 |
| | 5702821 | 48 | | 609419 | 62 | | | | | | | | 2066379 | 75 |
| | 5730763 | 45 | | 833785 | 54 | | | | | | | | 2107928 | 47 |
| | 5730765 | 45 | | 920012 | 53 | | | | | | | | 2176479 | 51 |

| | | | | | | | | | | | | | | |
|---|---|---|---|---|---|---|---|---|---|---|---|---|---|---|
| | 6148666 | 45 | | 949193 | 48 | | | | | | | | 2243380 | 68 |
| | 6372102 | 48 | | 1056161 | 53 | | | | | | | | 2263527 | 50 |
| | 6378983 | 48 | | 1127851 | 47 | | | | | | | | 2353268 | 69 |
| | 6601313 | 45 | | 1197559 | 58 | | | | | | | | 2600643 | 67 |
| | 6631579 | 48 | | 1379300 | 54 | | | | | | | | 2730944 | 59 |
| | 7047360 | 51 | | 1407873 | 46 | | | | | | | | 2933960 | 48 |
| | 7234658 | 50 | | 1445758 | 48 | | | | | | | | 2986244 | 46 |
| | 7456118 | 44 | | 1458752 | 57 | | | | | | | | 3045678 | 48 |
| *3* | 101298 | 53 | | 1603755 | 58 | | | | | | | | 3053646 | 49 |
| | 213205 | 44 | | 1608890 | 51 | | | | | | | | 3135817 | 72 |
| | 2961278 | 44 | | 1750472 | 57 | | | | | | | | 3282759 | 76 |
| | 3088951 | 46 | | 1910541 | 66 | | | | | | | | 3365274 | 47 |
| | 3137256 | 58 | | 2003959 | 45 | | | | | | | | 3488229 | 44 |
| | 3406894 | 49 | | 2035770 | 58 | | | | | | | | 3623739 | 60 |
| | 3444604 | 45 | | 2092912 | 66 | | | | | | | | 3627226 | 57 |
| | 3454364 | 47 | | 2113392 | 57 | | | | | | | | 3752974 | 57 |
| | 3513247 | 48 | | 2335024 | 51 | | | | | | | | 3967000 | 77 |
| | 3583055 | 49 | | 2392788 | 60 | | | | | | | | 4282464 | 87 |
| | 3898475 | 51 | | 2413594 | 45 | | | | | | | | 4530503 | 60 |
| | 3945255 | 45 | | 2453054 | 48 | | | | | | | | 4535587 | 55 |
| | 4224383 | 45 | | 2497491 | 60 | | | | | | | | 4554873 | 49 |
| | 4429811 | 44 | 3 | 187945 | 48 | | | | | | | | 4667661 | 91 |
| | 4471207 | 45 | | 204729 | 44 | | | | | | | | 4684252 | 48 |
| | 4543503 | 44 | | 281272 | 45 | | | | | | | | 4726766 | 59 |
| | 4649906 | 45 | | 451457 | 82 | | | | | | | | 4851483 | 56 |
| | 4840554 | 46 | | 718286 | 57 | | | | | | | | 4987854 | 48 |
| | 4921823 | 53 | | 823624 | 44 | | | | | | | | 5113526 | 45 |
| | 5174680 | 56 | | 893498 | 51 | | | | | | | | 5168462 | 52 |
| | 5643186 | 48 | | 1199792 | 45 | | | | | | | | 5261350 | 67 |
| | 5709725 | 55 | | 1242559 | 63 | | | | | | | | 5334003 | 48 |
| | 5837315 | 55 | | 1289687 | 49 | | | | | | | | 5771499 | 49 |
| | 6632858 | 48 | | 1308572 | 55 | | | | | | | | 5991689 | 46 |
| | 6734824 | 49 | 4 | 71574 | 63 | | | | | | | | 6001265 | 105 |
| | 6807453 | 48 | | 114527 | 71 | | | | | | | | 6215950 | 66 |
| | 6870955 | 47 | | 184355 | 64 | | | | | | | | 6320326 | 97 |
| | 7035322 | 57 | | 197421 | 53 | | | | | | | | 6467649 | 53 |
| *4* | 553361 | 50 | | 205545 | 45 | | | | | | | | 6520764 | 44 |
| | 603290 | 44 | | 225482 | 46 | | | | | | | | 6612449 | 57 |
| | 736928 | 46 | | 230302 | 47 | | | | | | | | 6688719 | 44 |
| | 739889 | 55 | | 235949 | 44 | | | | | | | | 6720946 | 48 |
| | 768623 | 49 | | 238163 | 46 | | | | | | | | 6732676 | 62 |
| | 780822 | 49 | | 267475 | 44 | | | | | | | | 6739254 | 68 |
| | 896029 | 52 | | 418300 | 48 | | | | | | | | 6775284 | 64 |
| | 1104029 | 45 | | 458856 | 59 | | | | | | | | 6784859 | 47 |
| | 1173895 | 56 | | 466766 | 61 | | | | | | | | 6869061 | 82 |
| | 1527369 | 57 | | 488398 | 70 | | | | | | | | 6938508 | 55 |
| | 1567474 | 44 | | 489907 | 50 | | | | | | | | 7046427 | 44 |

| | | | | | | | | | | | | | | |
|---|---|---|---|---|---|---|---|---|---|---|---|---|---|---|
| | 1784856 | 50 | | 496346 | 61 | | | | | | | | 7046430 | 45 |
| | 2026632 | 54 | | 573762 | 49 | | | | | | | | 7076632 | 88 |
| | 2068685 | 50 | | 718749 | 48 | | | | | | | | 7090589 | 57 |
| | 3385036 | 54 | | 743035 | 72 | | | | | | | | 7162432 | 45 |
| | 3407599 | 49 | | 797289 | 44 | | | | | | | | 7232503 | 46 |
| | 3466471 | 55 | | 865862 | 52 | | | | | | | | 7280011 | 51 |
| | 3729954 | 46 | | 875049 | 53 | | | | | | | | 7304673 | 47 |
| | 3875934 | 49 | | 926205 | 44 | | | | | | | | | |
| | 3918601 | 44 | | 988489 | 47 | | | | | | | | Total | 98 |
| | 4051648 | 53 | | 1023140 | 50 | | | | | | | | Extrapolated | 271 |
| | 4093371 | 45 | | 1064234 | 61 | | | | | | | | | |
| | 4135606 | 47 | | 1203811 | 49 | | | | | | | | | |
| | 4997387 | 46 | | 1278994 | 49 | | | | | | | | | |
| | 5011280 | 57 | 5 | 259107 | 53 | | | | | | | | | |
| | 5046326 | 51 | | 417398 | 45 | | | | | | | | | |
| | 5191272 | 52 | | 642904 | 77 | | | | | | | | | |
| *5* | 392351 | 47 | | 698360 | 65 | | | | | | | | | |
| | 804168 | 47 | | 837530 | 59 | | | | | | | | | |
| | 804170 | 47 | | 859134 | 52 | | | | | | | | | |
| | 1172419 | 44 | | 875111 | 45 | | | | | | | | | |
| | 1272736 | 50 | 6 | 11264 | 76 | | | | | | | | | |
| | 1283384 | 45 | | 72562 | 109 | | | | | | | | | |
| | 1835586 | 46 | | 371917 | 53 | | | | | | | | | |
| | 1888376 | 52 | | 391575 | 51 | | | | | | | | | |
| | 2001911 | 47 | | 541092 | 82 | | | | | | | | | |
| | 2344692 | 54 | | 581483 | 64 | | | | | | | | | |
| | 2427331 | 44 | | 667362 | 60 | | | | | | | | | |
| | 2502269 | 46 | | 827503 | 47 | | | | | | | | | |
| | 2926562 | 47 | | 848816 | 75 | | | | | | | | | |
| | 3631408 | 62 | 7 | 102254 | 48 | | | | | | | | | |
| | 3784634 | 48 | | 170299 | 45 | | | | | | | | | |
| | 4185592 | 54 | | 223212 | 64 | | | | | | | | | |
| | 4223708 | 46 | | 396780 | 51 | | | | | | | | | |
| *6* | 306678 | 55 | | 432674 | 65 | | | | | | | | | |
| | 589813 | 51 | | 450416 | 45 | | | | | | | | | |
| | 674719 | 60 | | 516576 | 57 | | | | | | | | | |
| | 1443458 | 62 | | 562750 | 56 | | | | | | | | | |
| | 1769064 | 44 | | 572982 | 49 | | | | | | | | | |
| | 1854512 | 49 | | 633540 | 51 | | | | | | | | | |
| | 2250217 | 48 | | 742582 | 44 | | | | | | | | | |
| | 2250221 | 48 | | 776609 | 66 | | | | | | | | | |
| | 2341266 | 43 | | | | | | | | | | | | |
| | 2344692 | 60 | | Total | 132 | | | | | | | | | |
| | 2426792 | 60 | | | | | | | | | | | | |
| | 2482603 | 50 | | | | | | | | | | | | |
| | 2927138 | 51 | | | | | | | | | | | | |
| | 2999567 | 44 | | | | | | | | | | | | |

| | | |
|---|---|---|
| | 3450386 | 57 |
| | 3529589 | 47 |
| | 3584584 | 47 |
| | 3698942 | 47 |
| | 3891647 | 50 |
| | 4371486 | 49 |
| | 4675850 | 45 |
| | 4757085 | 47 |
| | 4875608 | 57 |
| | 4879740 | 44 |
| | 5169078 | 50 |
| | 5298038 | 58 |
| | 5382903 | 48 |
| | 5751018 | 48 |
| *7* | 1272854 | 58 |
| | 1476943 | 64 |
| | 1500549 | 49 |
| | 1754112 | 58 |
| | 1862718 | 59 |
| | 1973673 | 47 |
| | 1997068 | 57 |
| | 2082703 | 47 |
| | 2133933 | 47 |
| | 2136350 | 53 |
| | 2973766 | 56 |
| | 3382630 | 53 |
| | Total | 176 |

| ***Saccharomycopsis fibuligera ACX*** | | | **Saccharomyces cerevisiae Y55** | | | **Aspergillus flavus NRRL3357** | | | **Neopestalotiopsis rosae 19-02** | | |
|---|---|---|---|---|---|---|---|---|---|---|---|
| *Chr* | Coor | bp | Chr: | Coor | bp | Chr | Coor | bp | Chr | Location | bp |
| *1* | 272788 | 45 | 1 | 0 | >44 bp | 3 | 592961 | 73 | 1 | 716909 | 53 |
| | 635588 | 49 | 2 | 778599 | 290 | 1 | 4815870 | 60 | | 1316839 | 60 |
| | 1029951 | 46 | 3 | 0 | NA | 1 | 5250063 | 60 | | 1869972 | 55 |
| | 1181878 | 44 | 4 | 4526 | 80 | 1 | 1959115 | 59 | | 3874500 | 45 |
| | 1373434 | 57 | 5 | 0 | NA | 2 | 1032759 | 59 | | 6041401 | 70 |
| | 1724098 | 55 | 6 | 277752 | 130 | 1 | 4079501 | 54 | | 6134764 | 61 |
| | 1944628 | 51 | 7 | 548397 | 46 | 2 | 3210941 | 53 | | 6991196 | 51 |
| | 2119444 | 48 | | 1022987 | 47 | 8 | 660991 | 52 | | 7015799 | 66 |
| | 2127913 | 53 | 8 | 0 | NA | 8 | 1342333 | 52 | 2 | 276182 | 82 |
| | 2304330 | 55 | 9 | 0 | NA | 3 | 4843136 | 49 | | 303144 | 67 |
| | 2345373 | 50 | 10 | 0 | NA | 8 | 3095989 | 49 | | 656060 | 53 |
| | 2495123 | 52 | 11 | 72146 | 58 | 8 | 1550755 | 48 | | 716187 | 49 |
| | 2551749 | 44 | | 648580 | 46 | 8 | 3010368 | 48 | | 1244574 | 62 |
| | 2653518 | 46 | 12 | 0 | NA | 1 | 3008144 | 47 | 3 | 468337 | 54 |
| | 2871342 | 46 | 13 | 0 | NA | 2 | 2426547 | 46 | | 1381802 | 49 |
| | 3471717 | 49 | 14 | 288055 | 48 | 3 | 2551031 | 46 | | 1440598 | 46 |

| | | | | | | | | | | | |
|---|---|---|---|---|---|---|---|---|---|---|---|
| *2* | 910788 | 58 | | 401996 | 51 | 5 | 3715764 | 45 | | 1528792 | 68 |
| | 1678228 | 44 | 15 | 922517 | 50 | 4 | 3535411 | 44 | | 1529957 | 74 |
| | 1771397 | 63 | 16 | 0 | NA | 5 | 3960053 | 44 | | 3945912 | 52 |
| | 2966517 | 55 | | Total | 11 | 5 | 4155120 | 44 | | 4145109 | 48 |
| | 2972032 | 55 | | | | 5 | 4373383 | 44 | | 4201114 | 45 |
| | 3906339 | 45 | | | | | | | | 4688881 | 44 |
| | 4035599 | 46 | | | | | Total | 21 | | 4910527 | 59 |
| *3* | 36844 | 44 | | | | | | | | 5008454 | 57 |
| | 1406670 | 52 | | | | | | | 4 | 1833177 | 72 |
| | 2630325 | 46 | | | | | | | | 4449191 | 53 |
| | 2919624 | 59 | | | | | | | | 6709765 | 67 |
| *4* | 291104 | 46 | | | | | | | 5 | 37168 | 48 |
| | 487186 | 48 | | | | | | | | 151725 | 48 |
| | 633652 | 56 | | | | | | | | 1406285 | 73 |
| | 668142 | 47 | | | | | | | | 1630461 | 70 |
| | 975087 | 65 | | | | | | | | 5191298 | 52 |
| | 1009816 | 48 | | | | | | | | 6449099 | 91 |
| | 1538420 | 44 | | | | | | | | 8044184 | 45 |
| | 1589627 | 69 | | | | | | | 6 | 407189 | 88 |
| | 2210541 | 82 | | | | | | | 7 | 437059 | 76 |
| | 2593397 | 54 | | | | | | | | 895431 | 59 |
| | 564996 | 62 | | | | | | | | 1365912 | 53 |
| | 922947 | 56 | | | | | | | | Total | 38 |
| | 1015641 | 50 | | | | | | | | | |
| | 71464 | 46 | | | | | | | | | |
| *6* | 327134 | 52 | | | | | | | | | |
| | 533322 | 57 | | | | | | | | | |
| *7* | 228201 | 71 | | | | | | | | | |
| | 685803 | 55 | | | | | | | | | |
| | 804996 | 52 | | | | | | | | | |
| | 899788 | 56 | | | | | | | | | |
| | Total | 47 | | | | | | | | | |

Table S3: Detailed information of data summarized in Figure 7 showing each species tested with listed chromosomes and coordinates in order of homopolymeric tract size

| Species | Strain | Sequence Methodology | Year | NCBI Designation |
|---|---|---|---|---|
| **Candidozyma auris** | B11809 | ONT | 2024 | GCF_003013715.1 |
| **Magnusiomyces capitatus** | LA-1 | PacBio Sequel<br>Illumina HiSeq | 2020 | GCA_029678045.1 |
| **Blastobotrys mucifer** | 17G1312 | PacBio Sequel II | 2024 | GCA_036871495.1 |
| **Saccharomycopsis fibuligera** | ACX0001 | PacBio | 2022 | GCA_023497445.1 |
| **Saccharomyces cerevisiae** | Y55 | PacBio Sequel | 2020 | GCA_903819135.2 |
| **Monascus purpureus** | KUPM5 | ONT<br>Illumina Novaseq | 2022 | GCA_025999795.1 |
| **Aspergillus fumigatus** | Afir974 | ONT<br>Illumina | 2023 | GCA_028752175.1 |
| **Aspergillus flavus** | NRRL 3357 | PacBio Revio | 2025 | This study |
| **Pyricularia oryzae** | Br48 | PacBio 2<br>Ilumina MiSeq, HiSeq | 2020 | GCA_036493215.1 |
| **Trichoderma reesei** | QM6a | PacBio Sequel | 2023 | GCA_028871515.1 |
| **Fusarium verticillioides** | ITEM10027 | ONT<br>Illumina HiSeq | 2023 | GCA_031360185.1 |
| **Neopestalotiopsis rosae** | 19-02 | ONT | 2024 | GCA_046115455.1 |
| **Nigrospora osmanthi** | Z1 | PacBio | 2023 | GCA_029489945.1 |

Table S4: Species and strains tested together with the method of sequencing, year of release and the NCBI designation of the assemblies.
